# A Stochastic Simulation-Based Approach to Inform Relapsing Mouse Model (RMM) Study Design for Non-Clinical Assessment of Tuberculosis

**DOI:** 10.1101/2025.07.18.665504

**Authors:** James Clary, Jessica K. Roberts, Debra Hanna, Alessia Tagliavini, Sylvie Sordello, Anna Upton, David Hermann, Alexander Berg

## Abstract

The development of new regimens to treat tuberculosis (TB), the disease caused by *Mycobacterium tuberculosis* (Mtb), is critical to improving patient outcomes and decreasing global infectious disease mortality. Early evaluation of candidate regimens in non-clinical models of TB, such as the relapsing mouse model (RMM), remains an important step in prioritizing the most efficacious regimens for further clinical evaluation. Although RMM studies may be informative, they are also animal-, labor-, and time-intensive to complete and represent significant investment in time and resources during non-clinical development. Given the strong pipeline of regimens in development, identification of “leaner” RMM studies may have a significant impact on resource utilization, and hence we compared alternative study designs with the goal of identifying study attributes that can be modified to improve resource use, particularly animal use. By simulating relapse outcomes from “virtual” studies (i.e., groups mice treated for selected durations with control and hypothetical anti-TB regimens) followed by model-based analysis of the simulated data, we were able to compare the “true” (input) values with model estimates of time to 95% cure probability (T_95_) and assess bias and precision of competing designs. Using this approach, we demonstrated that 28% fewer mice could be used in RMM studies while maintaining low bias and a precision for T_95_ estimation within +/− 1-2 weeks for most regimens. Therefore, it is expected that RMM studies based upon the alternative designs evaluated herein may be employed to promote improved animal stewardship while generating informative data for decision making.

## INTRODUCTION

Tuberculosis (TB), the disease caused by *Mycobacterium tuberculosis*, continues to remain a global health challenge affecting approximately 10.8 million individuals with an estimated mortality of approximately 1.25 million in 2023. While this is an improvement over previous years, which were heavily impacted by the COVID-19 pandemic, global progress in combating TB remains slow. Although WHO guidelines for drug-susceptible TB have been updated to allow for a 4-month daily regimen hi-dose Rifapentine, Moxifloxacin, Isoniazid and Pyrazinamide (RIPE), the previous standard of care regimen HRZE regimen (2-month combination of Isoniazid, Rifampin, Pyrazinamide, and Ethambutol followed by 4 months of Isoniazid and Rifampin) remains as first-line therapy.^[1]^ This highlights both the need for additional research into the development of new TB regimens as well as the potential for treatment shortening through new combination regimens, as the 4-month RIPE regimen can provide adequate treatment in a subset of patients while shortening treatment by 2 months.^[2]^ A treatment duration of 3 to 4 months or less total has been identified by the WHO as the minimal requirement for new regimens targeting drug-susceptible TB, with the optimal duration targeted at 2 months or less for the development of a pan-TB regimen (i.e., first-line therapy for any active TB, regardless of strain).^[3]^

A key consideration in the development of a shorter duration pan-TB regimen is that any such regimen is expected to include one or more novel drugs that are combined and dose-optimized for maximal efficacy and safety.^[3]^ Given the strong pipeline of new compounds, efficiently testing combinations early in development is critical to identify those that are most promising as candidate pan-TB regimens. Non-clinical studies, such as the BALB/c relapsing mouse model of pulmonary TB (RMM), are important tools to help inform potential regimen selection to expedite and prioritize promising agents for the clinic. The RMM has been extensively used to guide selection of candidate regimens for further development by evaluating by quantifying the proportion of treated mice exhibiting relapse following administration of selected drug combinations.^[4]^ Our recent work has demonstrated that the utility of the RMM in assessing relative regimen performance is greatly enhanced through the use of a model-based meta-analysis approach, whereby data from multiple studies is pooled and analyzed simultaneously. Specifically, we employed a mixed-effects logistic regression approach to analyzed data from 28 RMM studies to determine the treatment duration-dependent relapse probability for a given regimen.^[5]^ This method enabled the calculation of main metrics included treatment duration required to reach relapse probabilities of interest (e.g., time to 5% relapse probability, or inversely, the time to 95% probability of cure), while also quantifying and adjusting for the impact of study-level covariates on treatment response. Importantly, as the magnitude of inter-study variability was quantified, our approach allows for “apples-to-apples” comparison of all regimens across studies.

Although model-based meta-analysis is readily extendible to include emerging data from new and ongoing studies, an important caveat is that reliable estimation of treatment effects depends upon the size and informativeness of the underlying dataset. The ability of the model to estimate relative regimen performance relied heavily on data pooling to provide sufficient data for analysis. Even with data pooling, the available data for certain regimens (for example, only from a single treatment-duration) was too sparse to precisely estimate the key metrics of interest. These data limitations in part reflect that RMM studies have historically been designed to favor relatively large numbers of animals to enable statistical comparisons across groups at a limited set of treatment durations.^[6]^ Such studies, although statistically robust, were not designed to generate data that would be maximally informative for estimation of model-based parameters. Rather, when analyzed using a longitudinal model-based analysis, RMM studies should generate data suitable for accurate estimation of each regimen’s treatment duration vs. relapse probability curve. Key study attributes that may impact the data generated for estimation purposes include overall sample size, total mice per regimen, number and distribution of mice by treatment duration (timepoint), number of regimens arms in the study, inclusion of historical controls, range of regimen response (efficacy), and other study-level covariates that may influence treatment response (for example, inoculum size).^[5]^

To understand the relationships between study attributes and data informativeness for model-based analysis, we have performed an *in-silico* model-based evaluation of RMM study design. This builds upon our previous work by using the mixed-effects modeling framework to simulate the outcomes of virtual RMM studies and then “re-estimating” the simulated data using an updated model-based approach to generate measures of bias and precision for model-based parameters. By comparing the results of the simulation / re-estimation outputs across simulations, the impact of selected study attributes (i.e., number of mice per arm, number of mice per timepoint, and regimen selection / efficacy) were investigated to inform potential modifications to RMM study design. The overall goal was to identify study designs that would not only generate more informative data for model-based analysis, but would also remain logistically feasible, cost-effective, and promote minimal animal use.

## RESULTS

### Comparative performance of RMM study designs

#### Simulation Round 1

An initial round of simulations (Simulation Round 1) compared a baseline design with a set of proposed alternative designs and a high-performance benchmark (*viz*. “ultimate”) design (Table 1). All of the proposed designs, including the baseline design, already represent a departure from the typical historical design in that they allocate the total number of animals over a greater number of timepoints; for example, in the Baseline design, there are 6 mice assigned to each of 6 treatment durations (36 mice per arm) whereas in a historical design (see for example, Tasneen et al. ^[7]^), there would be half as many treatment durations with more than two times the number of mice evaluated at each. This was done purposefully to ensure that sufficient timepoints would be available to enable model-based estimation of the cure probability curve, and because such designs are already being implemented in practice.^[8]^ Twelve hypothetical regimens were simulated with intermediate efficacy relative to the two control arms (BPaMZ and HRZE) which were also included. Corresponding model parameter values for all regimens are displayed in Table 2 and the simulated (“true”) cure probability versus treatment duration curves are displayed in Figure 1. It should be noted that the regimens depicted in Figure 1 were fully hypothetical as they were not based on any regimens that had been studied as of the time of the analysis but were generated to represent plausible profiles for *de novo* regimens to be evaluated in future RMM studies.

**FIG 1.**
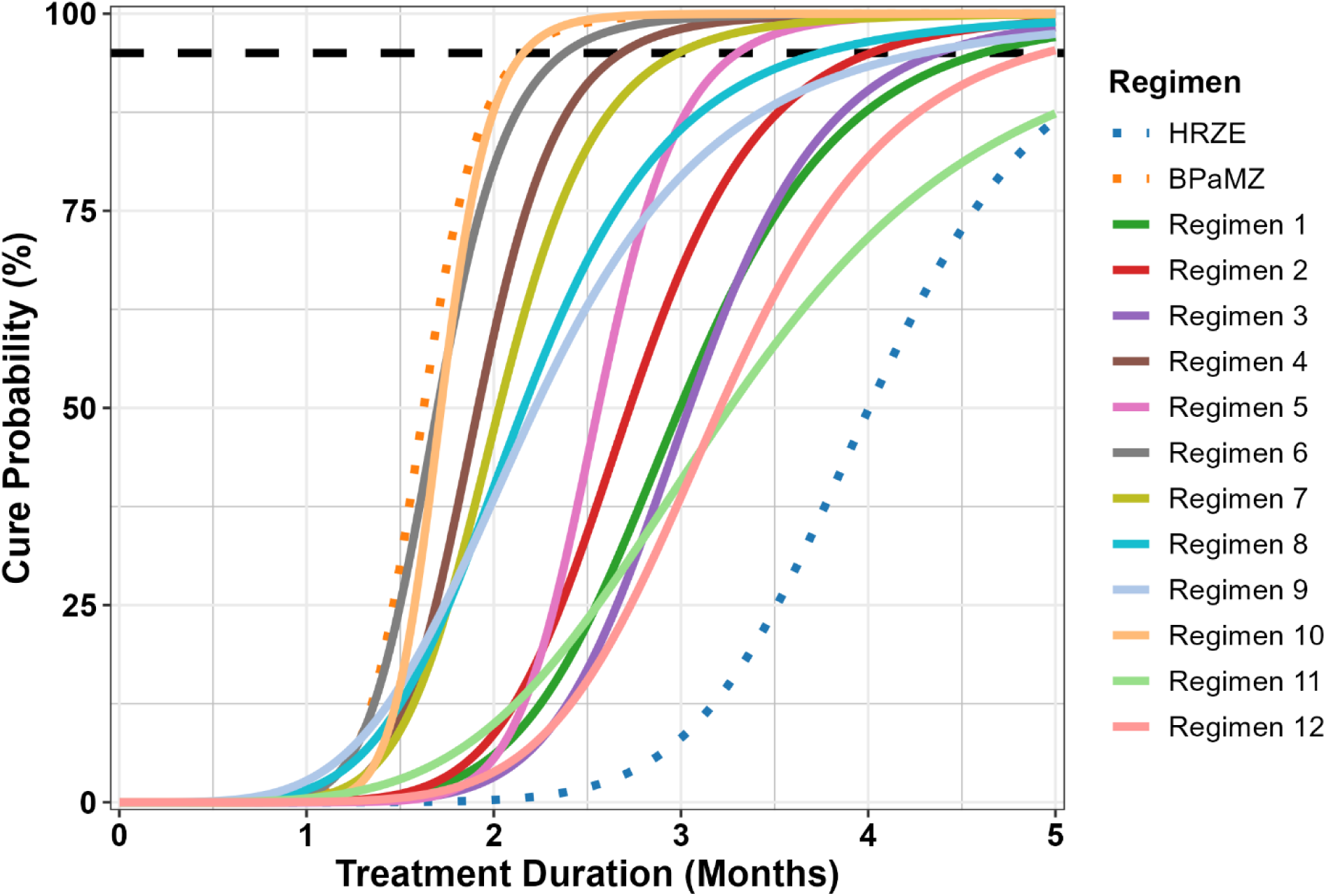
“True” cure probability versus treatment duration profiles for regimens used in Simulation Round 1. BPaMZ and HRZE (dotted lines) were control arms for the studies. All others were hypothetical regimens intended to represent plausible profiles. Dashed black line indicates 95% cure probability.

**TABLE 1:** Study Designs Evaluated – Round 1.

| Design | Description | Number of<br>Regimens | Treatment Durations<br>(months) | Mice Per<br>Duration | Total<br>Mice |
| --- | --- | --- | --- | --- | --- |
| Baseline | Original Design | 14 | 0.5, 1, 1.5, 2, 2.5, 3 | 6 | 504 |
| “Ultimate” | High performance benchmark | 14 | 0.5, 1, 1.5, 2, 2.5, 3, 3.5, 4, 4.5, 5 | 30 | 4200 |
| Proposed 1 | Omit two-week duration | 14 | 1, 1.5, 2, 2.5, 3 | 6 | 420 |
| Proposed 2 | Remove 3 study arms (HRZE,<br>Regimen 4, Regimen 11) | 11 | 0.5, 1, 1.5, 2, 2.5, 3 | 6 | 396 |
| Proposed 3 | Reduce to 5 mice per duration | 14 | 0.5, 1, 1.5, 2, 2.5, 3 | 5 | 420 |
| Proposed 4 | Reduce to 3 mice at two weeks<br>and 5 mice at other durations | 14 | 0.5, 1, 1.5, 2, 2.5, 3 | 5 | 392 |

**TABLE 2:**
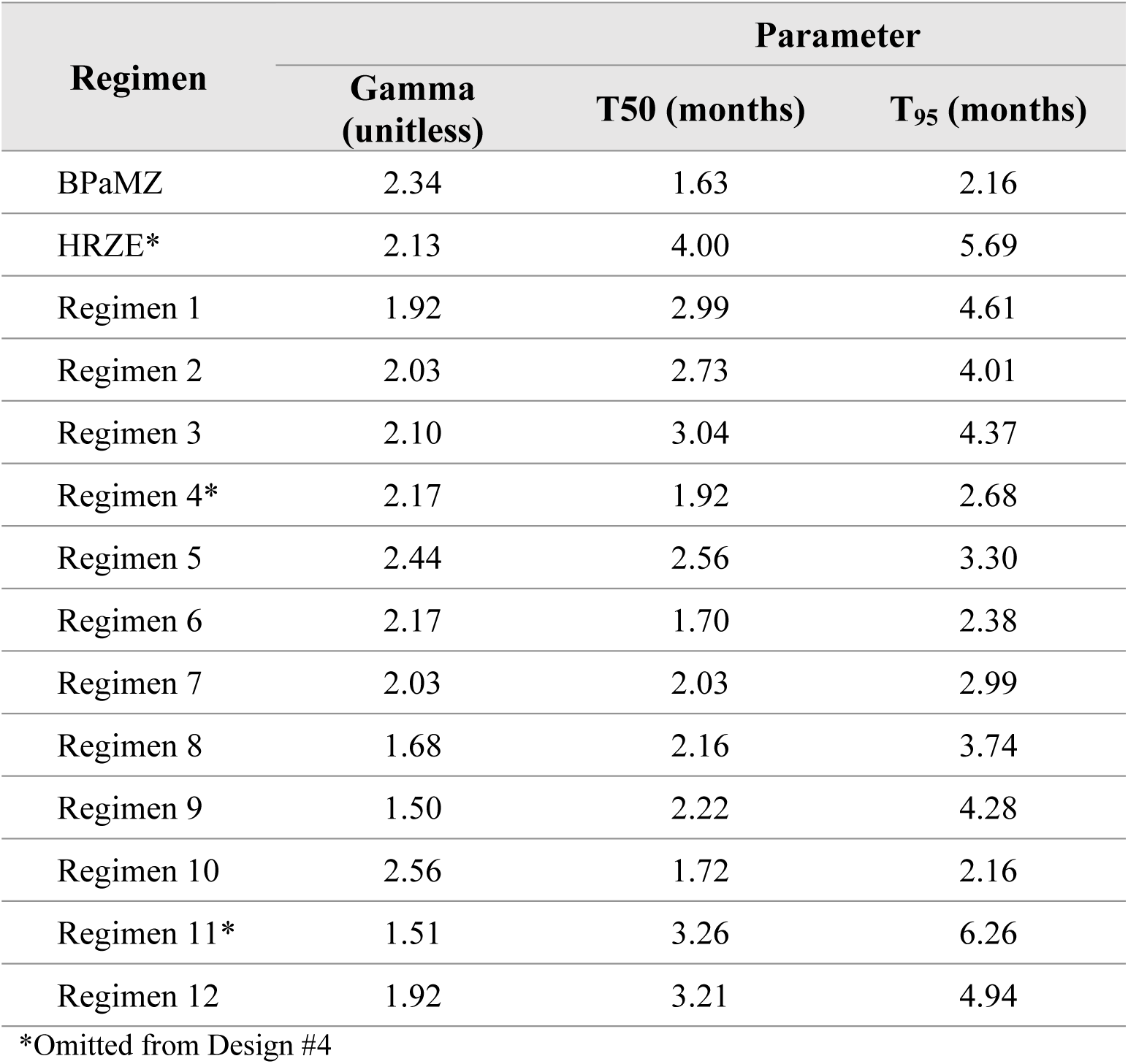
Hypothetical and Control Regimens – Simulation Round 1.

Overall, the baseline study design and proposed alternatives performed similarly across regimens, with no marked degradation in estimation of time to 95% probability of cure (T_95_, equivalent to the time to 5% probability of relapse) for any of the latter designs despite decreasing the number of mice by 17 to 22% or, in “Proposed 2”, also decreasing the number of regimens in the study by 3. In all cases, a median negative bias of up to 2 weeks was observed as was marked variability between regimens in the range of bias seen across simulations (Figure 2). This may also be seen in overlays of “true” relapse probability versus treatment duration profiles with the distribution of profiles estimated from the simulations (Supplementary Figures 3 through 16). The hypothetical regimens with the poorest accuracy and precision in T_95_ estimation were those that were the least efficacious; that is, those regimens where the steep part of the curve was mostly beyond the last time point (Regimens 1, 3, 11, and 12). These regimens exhibited median bias values in the range of –1 to –4 weeks and large interquartile ranges (IQR) across all study designs. In contrast, those regimens that exhibited the sharpest increase in cure probability (Regimens 4, 5, 6, 7, and 10) were relatively well estimated, with IQRs in the range of +/− 2 weeks.

**FIG 2.**
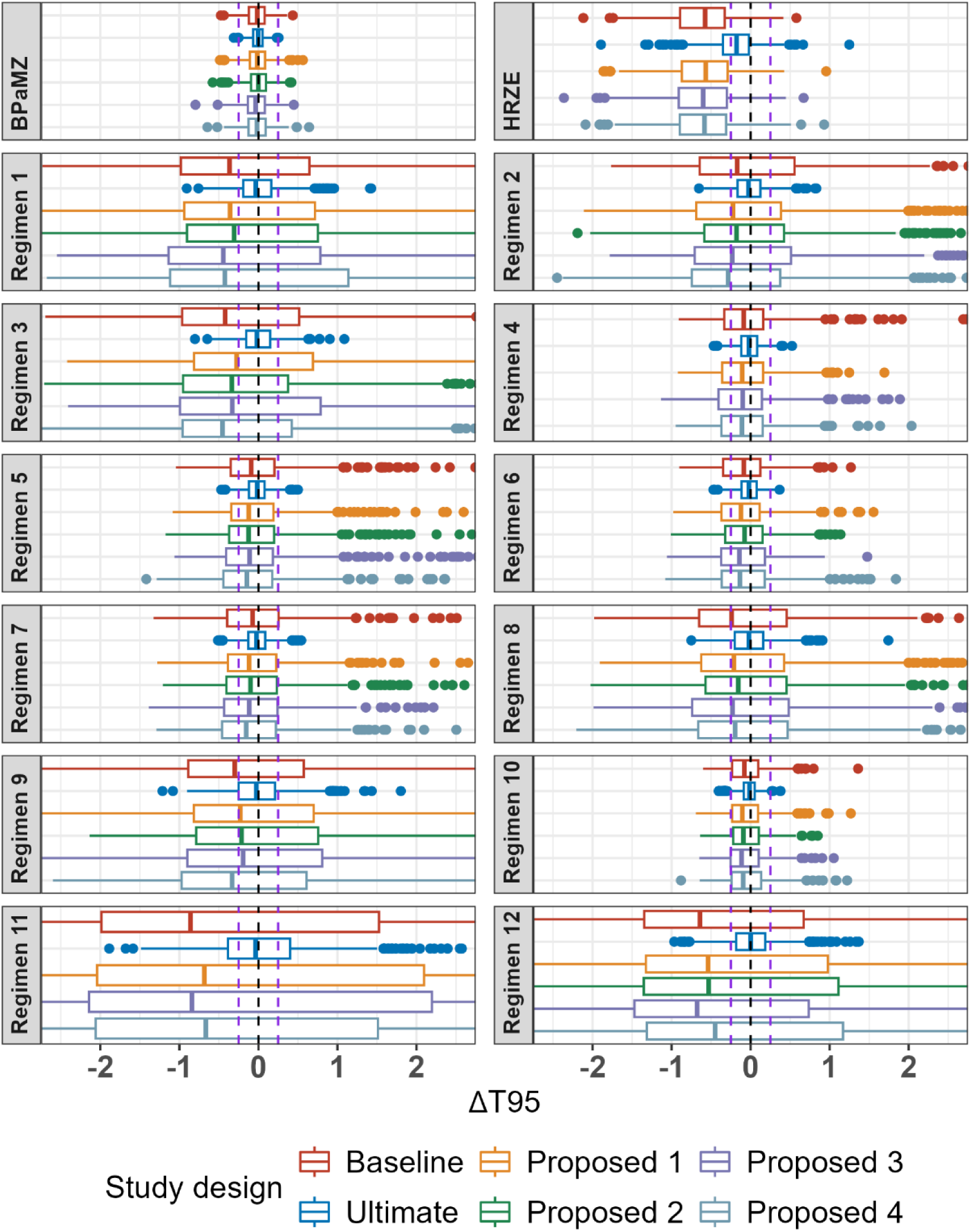
Bias plot of T_95_ (months) by regimen and study design from Simulation Round 1. Boxplots represent interquartile range, and lines represent 1.58* interquartile range (IQR); points are outliers outside this range. Lines represent 0 and ±0.25 months (±1 week) bias.

All study designs were able to reliably estimate the T_95_ value for the BPaMZ control arm, which was the most efficacious regimen with a similarly steep profile. Although the BPaMZ estimation was also supported by experimental data included in the analysis dataset during re-estimation, in the case of the HRZE control arm (ranked 13 out of the 14 regimens in terms of efficacy) including historical data was not sufficient to overcome the bias associated with having insufficient data points in the upper portion of the cure probability vs. treatment duration curve (*viz*., beyond 4 months). Further, omission of the HRZE control in “Proposed 2” did not result in worse performance vs. other designs which included both controls. This suggests that inclusion of more than one control arm may not be required to anchor the model to inform inter-study variability estimates, though it is acknowledged that within-study comparisons to the HRZE historical standard of care regimen may be of interest to provide confidence in study results.

Of note, while minimal differences were observed between the baseline and proposed alternatives, marked differences were observed when compared to the “ultimate” design. This implausibly large design, which included 4 additional treatment durations beyond 3 months and 5 to 6 times the number of mice per duration. Only the “ultimate” design was able to capture profiles where most of the curve is after 3 months, such as Regimen 12 (Supplementary Figure 16).

#### Simulation Round 2

The second round of simulations consisted of a focused assessment of baseline design versus two additional proposed alternative designs (“Proposed 5” and “Proposed 6”) that were selected based on the results from the first round of simulations (Table 3). As it was of specific interest to evaluate more potent anti-TB regimens in the RMM model (i.e, with similar performance to BPaMZ), a new set of hypothetical regimens were selected with T_95_ values between 1.75 and 3.5 months (Table 4 and Figure 3). It is noted that the parameters used for simulation of BPaMZ and HRZE profiles, while similar, were not identical between Simulation Rounds 1 and 2 (compare values in Tables 2 and 3). This reflects the iterative nature of the meta-analysis approach used for RMM data modeling, as data from additional studies featuring these regimens that became available were incorporated into the dataset and the model re-estimated between simulation rounds.

**FIG 3.**
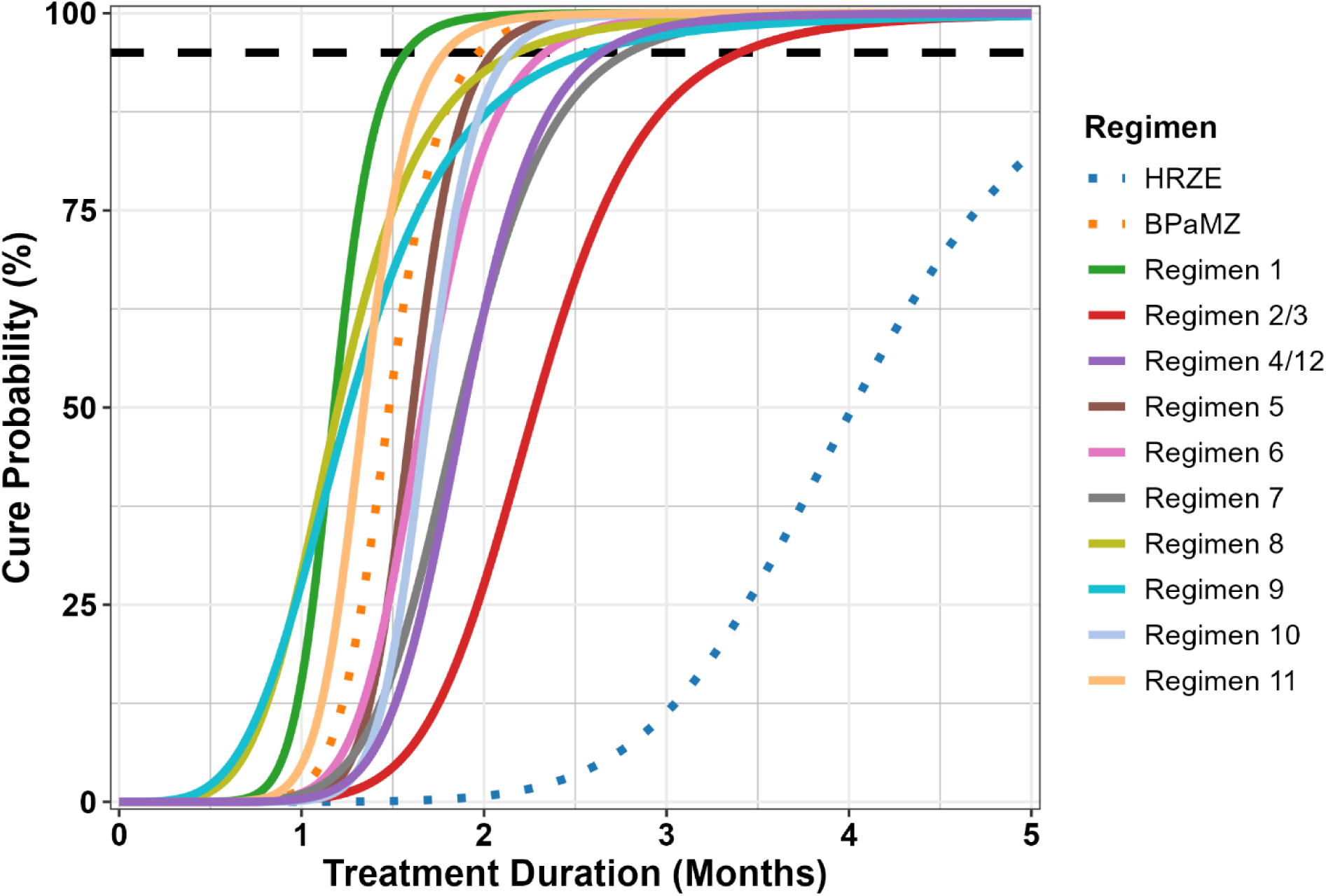
“True” cure probability versus treatment duration profiles for regimens used in Simulation Round 2. BPaMZ and HRZE (dotted lines) were control arms for the studies. All others were hypothetical regimens intended to represent plausible profiles. Dashed black line indicates 95% cure probability.

**TABLE 3:**
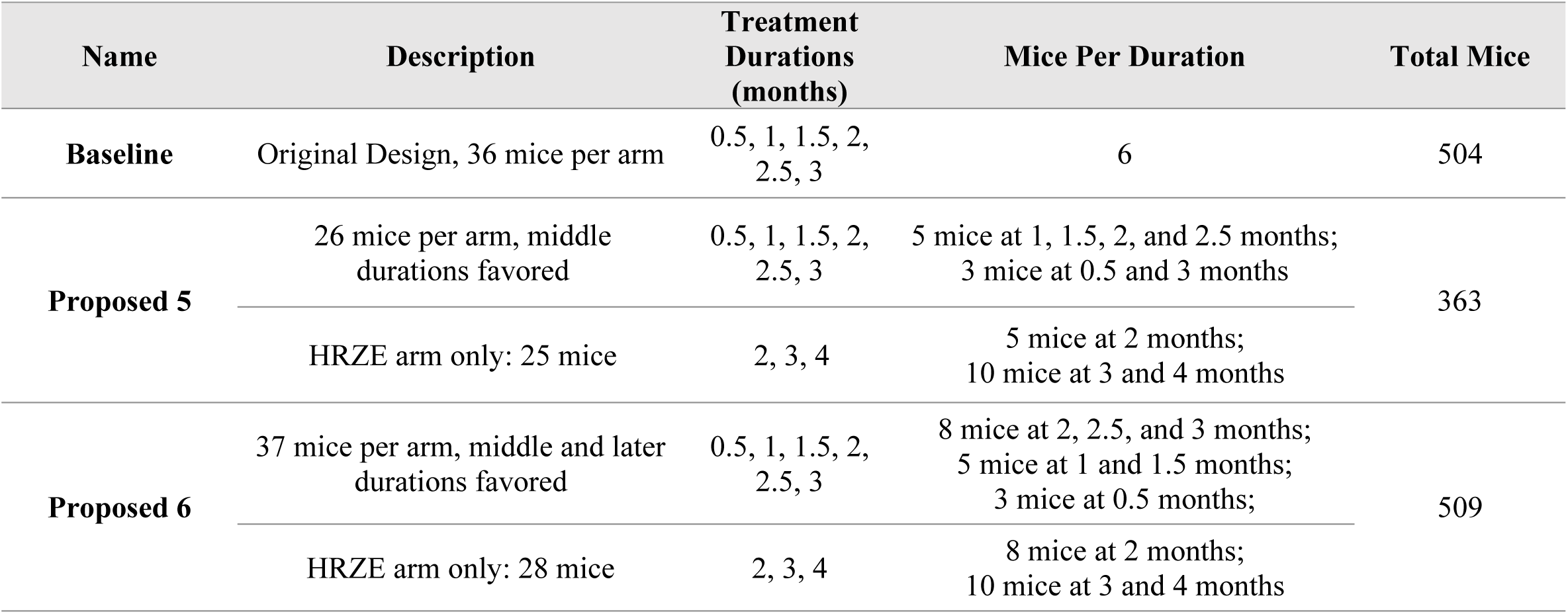
Study Designs Evaluated – Simulation Round 2.

| Name | Description | Treatment Durations (months) | Mice Per Duration | Total Mice |
| --- | --- | --- | --- | --- |
| **Baseline** | Original Design, 36 mice per arm | 0.5, 1, 1.5, 2, 2.5, 3 | 6 | 504 |
| **Proposed 5** | 26 mice per arm, middle durations favored | 0.5, 1, 1.5, 2, 2.5, 3 | 5 mice at 1, 1.5, 2, and 2.5 months;<br>3 mice at 0.5 and 3 months | 363 |
|  | HRZE arm only: 25 mice | 2, 3, 4 | 5 mice at 2 months;<br>10 mice at 3 and 4 months |  |
| **Proposed 6** | 37 mice per arm, middle and later durations favored | 0.5, 1, 1.5, 2, 2.5, 3 | 8 mice at 2, 2.5, and 3 months;<br>5 mice at 1 and 1.5 months;<br>3 mice at 0.5 months; | 509 |
|  | HRZE arm only: 28 mice | 2, 3, 4 | 8 mice at 2 months;<br>10 mice at 3 and 4 months |  |

**TABLE 4:** Hypothetical and Control Regimens – Simulation Round 2.

| Regimen | Parameter |  |  |
| --- | --- | --- | --- |
|  | Gamma<br>(unitless) | T50<br>(months) | T<sub>95</sub><br>(months) |
| BPaMZ | 2.03 | 1.48 | 1.97 |
| HRZE | 1.95 | 4.02 | 6.11 |
| Regimen 1 | 2.33 | 1.18 | 1.57 |
| Regimen 2 / 3 | 2.00 | 2.28 | 3.40 |
| Regimen 4 / 12 | 2.00 | 1.89 | 2.64 |
| Regimen 5 | 2.17 | 1.61 | 2.05 |
| Regimen 6 | 2.00 | 1.67 | 2.34 |
| Regimen 7 | 1.60 | 1.87 | 2.79 |
| Regimen 8 | 1.42 | 1.20 | 2.18 |
| Regimen 9 | 2.52 | 1.26 | 2.57 |
| Regimen 10 | 2.33 | 1.69 | 2.14 |
| Regimen 11 | 2.17 | 1.34 | 1.78 |

As in the first round of simulations, the original and proposed designs performed similarly across regimens for T_95_ estimation (Figure 4 and Table 5), although smaller bias and greater precision was observed for all hypothetical regimens in Simulation Round 2. These improvements are attributed in part to regimen and timepoint selection, as T_95_ values fell within the range of timepoints for all but two hypothetical regimens in second round. Additionally, the inclusion of data from additional studies, and a change in estimation approach from maximum likelihood estimation to Bayesian analysis using Markov Chain Monte Carlo (MCMC) in Simulation Round 2 also improved bias and precision. This is evidenced by the improvements seen for the BPaMZ and HRZE control arms when comparing the results for the Baseline design across simulation rounds (refer to Figures 2 and 4 for comparison). It is noted that the latter analysis approach is consistent with current methods employed by our group for RMM data analysis and illustrates how model-based approaches may be improved as analyses are repeatedly updated to incorporate and adapt to emerging data.

**FIG 4.**
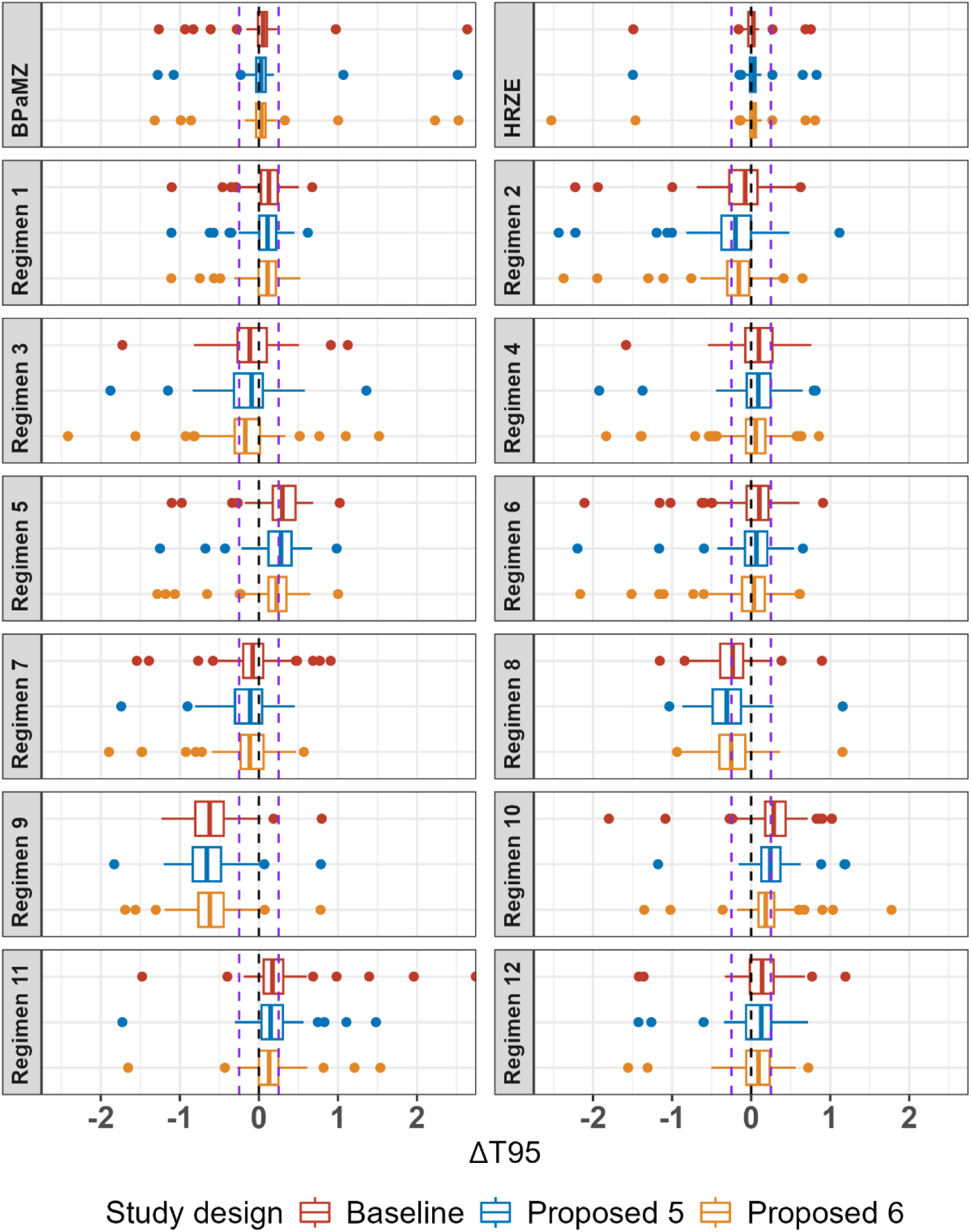
Bias plot of T_95_ (months) by regimen and study design from Simulation Round 2. Boxplots represent interquartile range, and lines represent 1.58* interquartile range (IQR); points are outliers outside this range. Lines represent 0 and ±0.25 months (±1 week) bias.

**TABLE 5:** Comparison of “true” T_95_ values in ascending rank order with model estimates from Simulation Round 2 (median and 5^th^ – 95^th^ percentiles)

| Regimen | “True” Values | | $T_{95}$ estimate (months) | | | $T_{95}$ Bias (months) | | |
| --- | --- | --- | --- | --- | --- | --- | --- | --- |
| | $T_{95}$ Rank Order | $T_{95}$ (months) | Baseline Design | Proposed Design 5 | Proposed Design 6 | Baseline Design | Proposed Design 5 | Proposed Design 6 |
| Regimen 1 | 1 | 1.57 | 1.70<br>(1.36, 1.97) | 1.68<br>(1.4, 1.92) | 1.68<br>(1.39, 1.92) | 0.13<br>(-0.21, 0.40) | 0.11<br>(-0.17, 0.35) | 0.11<br>(-0.18, 0.35) |
| Regimen 11 | 2 | 1.78 | 1.96<br>(1.71, 2.31) | 1.93<br>(1.63, 2.27) | 1.91<br>(1.66, 2.2) | 0.17<br>(-0.08, 0.53) | 0.15<br>(-0.15, 0.49) | 0.13<br>(-0.12, 0.42) |
| BPamZ | 3 | 1.97 | 2.02<br>(1.87, 2.13) | 2.00<br>(1.84, 2.12) | 2.00<br>(1.86, 2.14) | 0.05<br>(-0.09, 0.16) | 0.03<br>(-0.13, 0.15) | 0.03<br>(-0.11, 0.17) |
| Regimen 5 | 4 | 2.05 | 2.35<br>(1.97, 2.7) | 2.33<br>(1.95, 2.64) | 2.27<br>(1.94, 2.56) | 0.30<br>(-0.08, 0.65) | 0.28<br>(-0.10, 0.59) | 0.23<br>(-0.11, 0.52) |
| Regimen 10 | 5 | 2.14 | 2.43<br>(2.11, 2.77) | 2.39<br>(2.12, 2.67) | 2.33<br>(2.06, 2.65) | 0.29<br>(-0.03, 0.63) | 0.24<br>(-0.02, 0.53) | 0.19<br>(-0.08, 0.50) |
| Regimen 8 | 6 | 2.18 | 1.95<br>(1.59, 2.32) | 1.87<br>(1.49, 2.28) | 1.93<br>(1.51, 2.34) | -0.23<br>(-0.58, 0.14) | -0.31<br>(-0.68, 0.11) | -0.25<br>(-0.67, 0.16) |
| Regimen 6 | 7 | 2.34 | 2.44<br>(2.00, 2.72) | 2.4<br>(1.96, 2.71) | 2.37<br>(1.95, 2.78) | 0.10<br>(-0.33, 0.39) | 0.07<br>(-0.38, 0.38) | 0.04<br>(-0.39, 0.44) |
| Regimen 9 | 8 | 2.57 | 1.95<br>(1.53, 2.39) | 1.91<br>(1.46, 2.41) | 1.95<br>(1.57, 2.36) | -0.62<br>(-1.04, -0.18) | -0.66<br>(-1.11, -0.16) | -0.62<br>(-1.00, -0.21) |
| Regimen 4 | 9/10 | 2.64 | 2.74<br>(2.40, 3.12) | 2.74<br>(2.35, 3.13) | 2.7<br>(2.24, 3.00) | 0.10<br>(-0.25, 0.47) | 0.09<br>(-0.29, 0.48) | 0.06<br>(-0.40, 0.35) |
| Regimen 12 | 9/10 | 2.64 | 2.78<br>(2.44, 3.24) | 2.77<br>(2.35, 3.18) | 2.74<br>(2.36, 3.09) | 0.14<br>(-0.20, 0.59) | 0.13<br>(-0.29, 0.53) | 0.09<br>(-0.28, 0.44) |
| Regimen 7 | 11 | 2.79 | 2.71<br>(2.29, 3.14) | 2.68<br>(2.30, 3.03) | 2.68<br>(2.27, 3.03) | -0.08<br>(-0.50, 0.35) | -0.11<br>(-0.49, 0.24) | -0.11<br>(-0.52, 0.24) |
| Regimen 2 | 12/13 | 3.40 | 3.33<br>(2.89, 3.77) | 3.20<br>(2.70, 3.73) | 3.25<br>(2.88, 3.63) | -0.07<br>(-0.51, 0.36) | -0.20<br>(-0.70, 0.33) | -0.15<br>(-0.53, 0.23) |
| Regimen 3 | 12/13 | 3.40 | 3.29<br>(2.90, 3.77) | 3.31<br>(2.83, 3.69) | 3.23<br>(2.85, 3.64) | -0.11<br>(-0.51, 0.37) | -0.09<br>(-0.57, 0.29) | -0.17<br>(-0.55, 0.23) |
| HRZE | 14 | 6.11 | 6.14<br>(6.05, 6.19) | 6.14<br>(6.05, 6.21) | 6.15<br>(6.04, 6.21) | 0.03<br>(-0.06, 0.08) | 0.03<br>(-0.06, 0.10) | 0.04<br>(-0.07, 0.10) |

In Simulation Round 2, most regimens showed a median bias within +/− 1 week with an interquartile range of approximately 1 week. Regimens with a bias outside this range included Regimens 5 and 10, which showed a median positive bias at or above +1 week for Baseline and Proposed 5 designs. These regimens exhibited similar T_95_ values of approximately 2.1 months and “steep” profiles as indicated by the small difference between T_95_ and the midpoint of the curve (given by the T50 estimate: 0.44 and 0.45 months for Regimen 5 and Regimen 10, respectively). Other “steep” hypothetical regimens that had a less than a 0.5-month difference between the midpoint and T_95_ included Regimens 1 and 11, which also showed a small positive bias. In contrast, Regimen 9, which was the “shallowest” hypothetical regimen (T_95_ = 2.57 months, time between T_50_ and T_95_ = 1.31 months), had a negative bias between –2 and –3 weeks and an IQR completely falling below –1.5 weeks for all study designs. While this suggests that both the shape of the profile and the overall efficacy may influence the direction and magnitude of bias, given that the IQR is narrow, in general the overall bias in model estimates remains small relative to the required treatment duration to achieve high cure rates (i.e., median absolute bias ranging from 0.03 to 0.66 months vs. T_95_ values of 1.6 to 3.4 months). When viewed as a Forest plot, a typical way of comparing regimen performance (Figure 5), there is generally good agreement between the “true” value and model estimates, with the former generally lying within the 5^th^ and 95^th^ percentiles of T_95_ values estimated across replicates. Although some differences in T_95_-based regimen rank order are indicated (Supplementary Table 1), this was expected given that T_95_ values for the most efficacious hypothetical regimens differ only on the order of days, whereas bias and precision estimates are on the order of weeks. For example, those that were the most incorrectly ranked (i.e., had a median rank that was more than one position different from their “true” rank), Regimens 5, 8, 9, and 10, all had “true” T_95_ values within an approximately 2-week range. In this situation, rank order assessment is challenging and can be particularly misleading as minor differences in estimation can result in ranking errors even though the magnitude of the estimation bias is small. However, as differences between “true” and observed rank order were seen with all proposed study designs, the simulations do not suggest a clear advantage of one design vs. another in terms of regimen rank order assessment.

**FIG 5.**
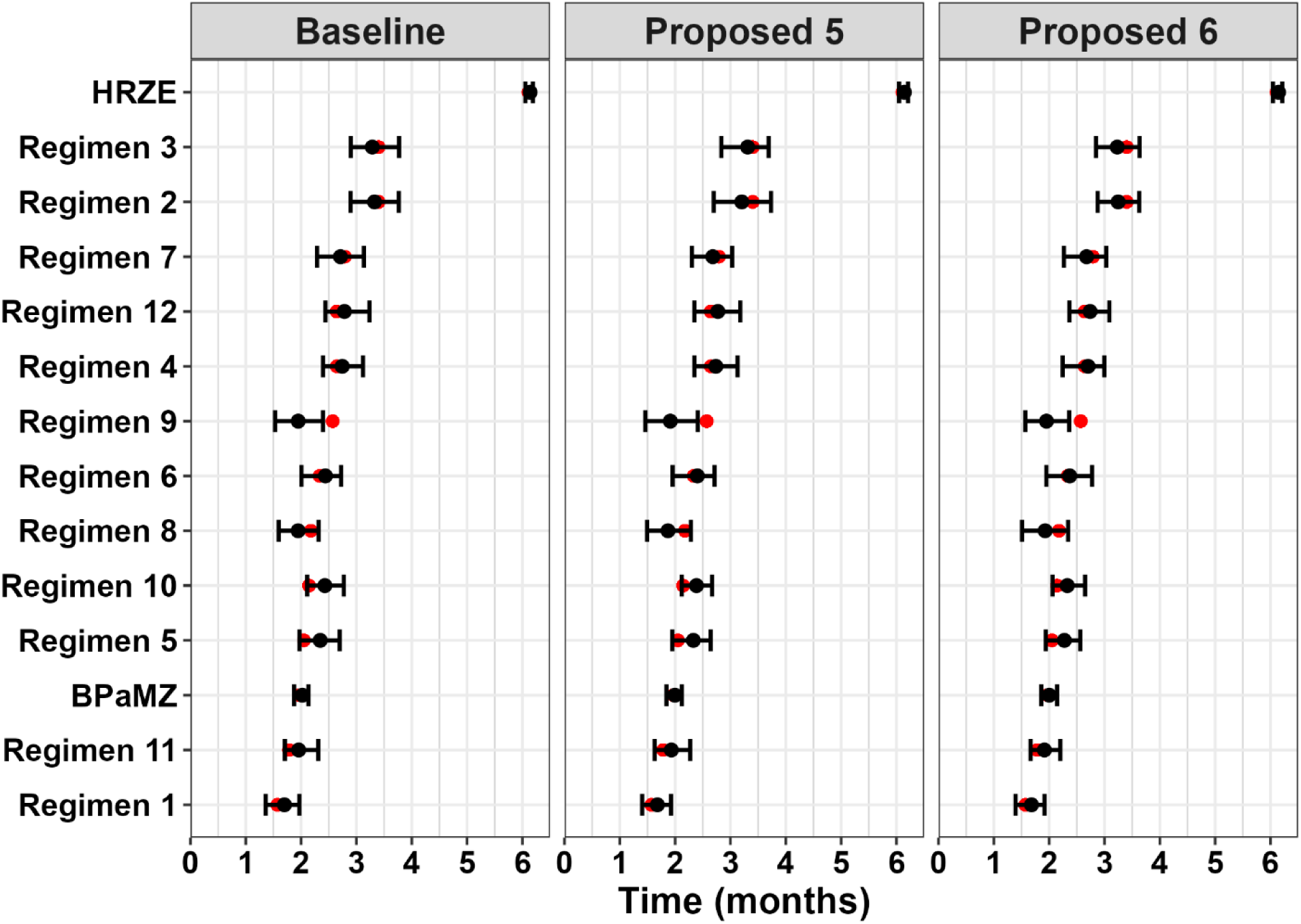
Comparison of “true” and estimated T_95_ values by regimen and study design from Simulation Round 2. Regimens are organized in descending rank order from least to most efficacious regimen based on the “true” value (red point). The black point (range) represents the median (5^th^ and 95^th^ percentiles) of T_95_ estimates across simulation replicates.

## DISCUSSION

By employing a simulation / re-estimation approach, we were able to compare RMM study designs to evaluate their relative performance in the reliable estimation of key metrics of interest (*viz*., T_95_). In general, the original and proposed alternative designs evaluated in this simulation study showed similar performance to each other for hypothetical and control regimens, with low bias for most regimens and a precision for T_95_ estimation within +/− 1 to 2 weeks. This is despite differences in the total number of mice, timepoints, mice per timepoint, and/or number of regimens. The only significant improvements in the simulation study were seen with the “ultimate” design (N = 4200 mice) included in the first simulation round, a completely unrealistic design which was included to benchmark the “best” possible performance. While the results for this design suggested that that further improvements may be possible for regimens where the T_95_ value lies beyond the range of treatment durations evaluated in the study, it is noted that such regimens would likely be less efficacious and therefore considered as lower priority. This obviates the return on the investment in more treatment durations and/or mice per treatment duration, especially to the level indicated with such an implausibly large study design. Of the other designs, although some improvement was suggested in the second simulation round with “Proposed 6” (N = 509 mice, similar in size to the Original Design) showing smaller bias for “steep” regimens (that is, Regimens 5 and 10), the improvement was minor as compared to “Proposed 5” (N = 363 mice, the smallest design evaluated). This is important in that animal stewardship continues to be important criteria in the development of research programs^[9,10]^, especially when large numbers of animals are planned for terminal sacrifice in critical non-clinical experiments such as the RMM. In this simulation study, we demonstrated that by using a model-based analysis approach and tuning the allocation of mice across more informative timepoints, a large 28% reduction in the number of mice (i.e., Proposed Design 5 versus the Baseline Design) can be achieved with minimal impact on T_95_ estimation. Moreover, these and additional (unpublished) simulations indicate that further reduction in animal usage could be achieved through other selected modifications (e.g., decreasing the numbers allocated to control arms or dropping one of the control arms from the design altogether). Taken as a whole, the results presented herein demonstrate that “leaner” RMM study designs that are more cost-effective and potentially more logistically feasible can significantly decrease animal use while remaining highly informative for decision making.

A key assumption in this comparative assessment is that all RMM studies conducted using the designs outlined herein would be analyzed using a longitudinal model-based approach. This methodology “fits” the relapse proportions at each treatment duration into a smooth curve for each regimen and treats the entire study as an experimental group for analysis. Such regression-based approaches are focused on obtaining reliable parameter estimates for cross-regimen comparison as compared to historical methods of analysis focused on statistical analysis to determine statistically significant differences in relapse proportions at specific timepoints. As the latter approach requires significantly larger sample sizes to obtain statistical significance^[6]^, the proposed study designs would not be sufficiently powered for such analyses.

Another consideration is that the specific approach utilized by our team is a model-based meta-analysis, which utilizes historical RMM study data to anchor model estimates and uses data from control arms as a bridge between the new and previous studies. The benefits of this method are that large datasets help to improve parameter estimates for fixed effects (i.e., treatment regimen and covariate parameters) and random effects (i.e., inter-study variability parameters), and that emerging data is analyzed in the context of previous RMM studies, thereby allowing for cross-study comparisons. This represents a potential limitation, as it is acknowledged that access to RMM study datasets may not be universal, and it is unlikely that datasets containing all relevant data will be available on an ongoing basis without significant investment in data management and data sharing (although it is noted that several RMM studies included in this and previous work^[5]^ are accessible via the TB-Platform for the Aggregation of Preclinical Experiments Data [TB-APEX] database).^[11]^ To address this limitation, rather than rely solely on historical data to stabilize parameter estimates, the MCMC Bayesian analysis methodology employed in Simulation Round 2 has the advantage of allowing the incorporation of prior information related to model parameter distributions. Although in this study non-informative “priors” were used for the hypothetical regimens, relevant information regarding likely regimen performance (i.e., parameter values), covariate effects, or random effects, can be incorporated using “informative” priors and therefore help to overcome data limitations. This advantage, as well as the multiple technical improvements observed in this study (for example, better parameter estimation, shorter run times, and improved model stability), has led to the adoption of MCMC as our standard approach for analysis of emerging RMM study data.

In summary, using a simulation-based approach, we were able to demonstrate that alternative RMM study designs were able to produce similar performance in calculating metrics of interest from a model-based analysis. By adjusting key design elements, including mice per treatment duration and the total number of treatment durations evaluated, one proposed study design (Proposed Design 5) was able to reduce the total number of mice by 28% while still maintaining good precision compared to the baseline design. This study design has since been implemented for multiple ongoing (unpublished results) and completed RMM studies, including that reported by Sordello et al.^[8]^ to successfully balance improved animal stewardship and while providing informative data to support the non-clinical evaluation of new and novel regimens.

## METHODS

### Statistical Model

The statistical model used for estimation purposes in this simulation / re-estimation study was an extension of the model of Berg et al.^[5]^ as implemented by Sordello et al ^[8]^ The updated model featured an inverse E_max_-type structure defined by two key parameters, midpoint (T50) and shape (γ). These parameters were estimated separately by regimen to define the typical relapse probability vs. treatment duration profiles for each regimen after adjusting for covariate effects and inter-study variability. The general model, which also includes the effect of inoculum amount (INOC) as a covariate, is described by the following Equations:

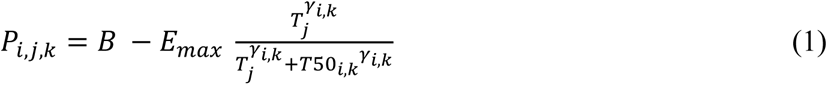

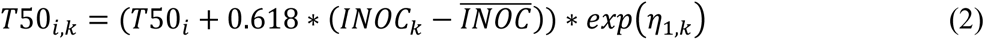

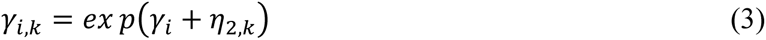

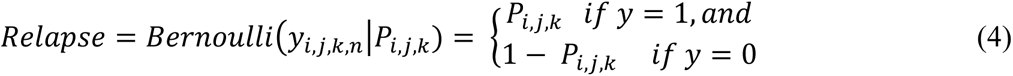

Where:

- *P_i,j,k_* is the probability of relapse for the i^th^ regimen at the j^th^ month in the k^th^ study;
- *B* is the baseline probability of relapse prior to treatment (Fixed to 1, assumes 0% cure with no treatment);
- *E_max_* is the maximum effect for all regimens (Fixed to 1, assuming 100% cure with continued treatment);
- *T_j_* is the j^th^ treatment duration (months);
- *T50_i,k_* is the midpoint of the curve, the treatment duration to achieve 50% relapse or cure probability for the i^th^ treatment in the k^th^ study;
- *γ_ik_* is the shape parameter (Hill coefficient) for the i^th^ treatment in the k^th^ study;
- *INOC_k_* is the inoculum covariate (log_10_ CFU) for the k^th^ study;
- 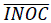 is the median inoculum value across all studies, corresponding to values of 3.65 and 4.01 log_10_ CFU for simulation rounds 1 and 2, respectively;
- η_1,*k*_ and η_2,*k*_ are random effects of the *k*^th^ study for *T_50_* and *γ*, respectively, assumed to be N(0,σ^2^) with an unstructured covariance matrix; and
- *y_i,j,k,n_* is the relapse status (1 = relapse, 0 = cure) of the *n*^th^ mouse in the *k*^th^ study receiving the *i*^th^ treatment at month *j*.

### Simulations

Simulations were performed to compare the baseline study design relative to proposed alternative designs that were logistically feasible, highly informative, cost-effective, and minimized animal use. Simulations were performed iteratively in two rounds, with the parameters evaluated in the second simulation round informed by results from the first simulation round. Hypothetical regimens (defined by T50 and γ values) were simulated to represent a range of plausible cure probability versus treatment duration profiles for *de novo* regimens. All simulations were performed in R v.4.0.3 as implemented via RStudio Workbench v1.4.1717-3.^[12]^ For each replicate (equivalent to one virtual RMM study), η estimates for T_50_ and γ were generated from the variance-covariance matrix using the *MASS* package^[13]^ (first round) or were sampled with replacement from study-specific values obtained from previous RMM study analyses (second round; unpublished data). The resulting η values were input to the model equations along with prespecified T50 and γ for the selected hypothetical or control regimens and an inoculum covariate value of 4.5 log_10_ CFU (representative of the inoculum used for planned studies). The corresponding study-, covariate-, and regimen-specific relapse probabilities at specific timepoints (treatment durations) were then used to simulate an outcome (relapse status; 0 or 1 for cure or relapse, respectively) for each “virtual” mouse in the study by drawing from a binomial distribution.

#### Simulation Round 1

Twelve hypothetical anti-TB regimens designs were used to assess a range of sterilization rates considered plausible for the unknown drug regimens. Two existing drug regimens (BPaMZ [Bedaquiline, Pretomanid, Moxifloxacin, and Pyrazinamide] and HRZE) were included as control regimens. Refer to Table 2 for the corresponding model parameters used for simulation of the various regimens. Six different study designs were assessed, including a baseline design corresponding to a real RMM study pending execution at the time, an “ultimate” design featuring implausibly high numbers of animals and timepoints (included to estimate the best possible performance characteristics), and four proposed alternative designs (namely Proposed 1 through 4). As preliminary simulations suggested that model convergence upon re-estimation could be low in certain simulation scenarios, a total of 1000 replicates were simulated to ensure adequate numbers of studies with reliable parameter estimates.

#### Simulation Round 2

A separate set of twelve hypothetical anti-TB drug regimens were generated to compare the baseline design with two additional proposed alternative study designs (Proposed 5 and 6). BPaMZ and HRZE were retained as controls arms, whereas two sets of regimens were duplicated (Regimens 2 / 3 and Regimens 4 / 12) to investigate the ability for the model to identify regimens with identical performance within the same study. Refer to Table 4 for the corresponding model parameters used for simulation of the various regimens. For each design, 200 replicates were simulated.

### Model re-estimation from simulated data sets

Each simulated replicate dataset was combined with available experimental RMM data to generate an estimation-ready dataset for each replicate. This was done to match how *de novo* study data is typically analyzed in practice by our group. The model was then re-estimated separately for each simulation iteration following addition of regimen specific T50 and γ parameters for the hypothetical regimens.

In the first simulation round, re-estimation was performed in NONMEM v7.3, with only models that achieved successful convergence included in the outputs. In the second simulation round, a Bayesian estimation approach (MCMC) was utilized using the statistical analysis software, Stan, as implemented by the RStan package in R.^[14]^ For MCMC analysis, normal distributions were used as prior for all model parameters. Weak priors were used for hypothetical regimen parameters (T50: mean = 1 to 2 months, standard deviation = 0.5 months; γ: mean = 2.3, standard deviation = 1), with informative priors for all other parameters (i.e., control regimen T50 and γ, inoculum covariate effect, and sigma [eta variance] parameters) based on mean and standard deviation estimates from the pooled meta-analysis dataset.

For reference, visual predictive checks stratified by RMM study index are provided in Supplemental Figures 1 and 2, which show model performance relative to the historical data for the models used simulation rounds 1 and 2, respectively, and demonstrate sufficient performance for estimation and simulation purposes.

### Comparison of Designs

Following re-estimation, T_95_ (time to 95% probability of cure) values were calculated for each replicate and each individual regimen, using Equation 5:

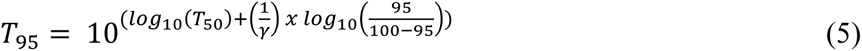

These metrics were summarized by design and compared to the true simulation to assess bias and overall predictive performance of the design. Bias for each simulation replicate was calculated as shown in Equation 6.

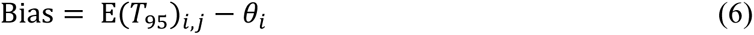

Where *E(T_95_)*_i,j_ is the estimated T_95_ value obtained from a given simulation replicate for each regimen, *i*, and study design, *j*, with *θ*_i_ denoting the “true” (input) T_95_ value for each simulated regimen. For each replicate, estimated cure probability versus treatment duration curves were simulated from the model estimates to obtain a distribution for each study design to graphically compare model estimates with the true values. Finally, each replicate’s T_95_ values were ranked, and the order was compared to the value used for simulations across designs to assess the ability of each design to differentiate regimens. All graphical representations were generated in R using the *ggplot2* package.^[15]^

### Data Availability

This report details the results of a simulation study and no experimental data was generated. Simulation code for generating simulated datasets is available upon request. Historical RMM study data used for re-estimation purposes is available to qualified researchers via the TB-APEX data platform available at https://c-path.org/tools-platforms/tb-apex/.

## ACKNOWLEDGMENTS

The authors acknowledge the contributions of Johns Hopkins University and Colorado State University in providing the relapsing mouse model datasets used in model development.

Proofreading and formatting support was provided by Rachel Ham (Allucent [US], LLC) in preparation of this manuscript. Project management support for this project provided by Lindsay Lehmann (Simulations Plus, Inc).

This work was supported by the Bill and Melinda Gates Foundation, which provided funding for the modeling effort to Simulations Plus, Inc and Allucent for manuscript preparation.

We declare no conflict of interest.

## APPENDIX

**STABLE 1:** Comparison of true and model-estimated regimen T_95_ rank order in ascending order – Median (5^th^-95^th^ percentiles) from Simulation Round 2.

| Regimen | “True” Rank | Original Design | Proposed Design 5 | Proposed Design 6 |
| --- | --- | --- | --- | --- |
| Regimen 1 | 1 | 1<br>(1, 3) | 1<br>(1, 3) | 1<br>(1, 4) |
| Regimen 11 | 2 | 3<br>(2, 6) | 3<br>(1, 6) | 3<br>(2, 6) |
| BPaMZ | 3 | 4<br>(2, 5) | 4<br>(2, 5) | 4<br>(2, 6) |
| Regimen 5 | 4 | 7<br>(4, 9) | 7<br>(4, 9) | 6<br>(4, 9) |
| Regimen 10 | 5 | 7<br>(6, 10) | 7<br>(5, 9) | 7<br>(5, 9) |
| Regimen 8 | 6 | 3<br>(1, 6) | 3<br>(1, 6) | 3<br>(1, 7) |
| Regimen 6 | 7 | 7<br>(5, 10) | 7<br>(4, 10) | 7<br>(4, 10) |
| Regimen 9 | 8 | 3<br>(1, 7) | 3<br>(1, 7) | 4<br>(1, 8) |
| Regimen 4 | 9/10 | 10<br>(7, 12) | 10<br>(7, 12) | 10<br>(8, 11) |
| Regimen 12 |  | 10<br>(8, 12) | 10<br>(8, 12) | 10<br>(8, 11) |
| Regimen 7 | 11 | 10<br>(7, 11) | 9<br>(7, 11) | 10<br>(7, 11) |
| Regimen 3 | 12/13 | 12<br>(11, 13) | 13<br>(11, 13) | 12<br>(11, 13) |
| Regimen 2 |  | 12<br>(11, 13) | 12<br>(10, 13) | 12<br>(11, 13) |
| HRZE | 14 | 14<br>(14, 14) | 14<br>(14, 14) | 14<br>(14, 14) |

**SFIG1.**
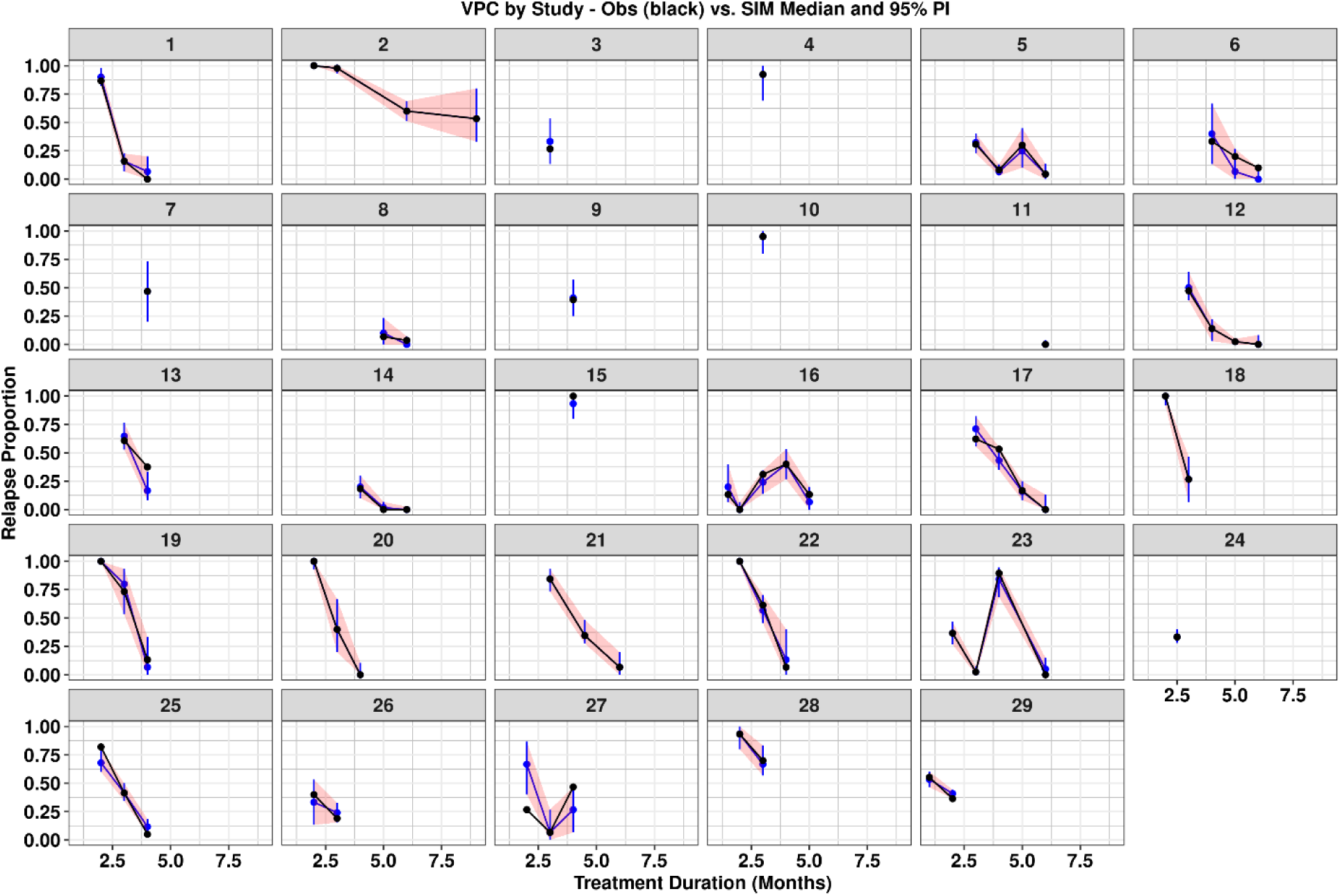
Visual predictive check (VPC) for the model used in Round 1 simulations. Red shaded regions and blue error bars represent the 90% Prediction Interval for the model. Black points and black lines represent the actual data.

**SFIG2.**
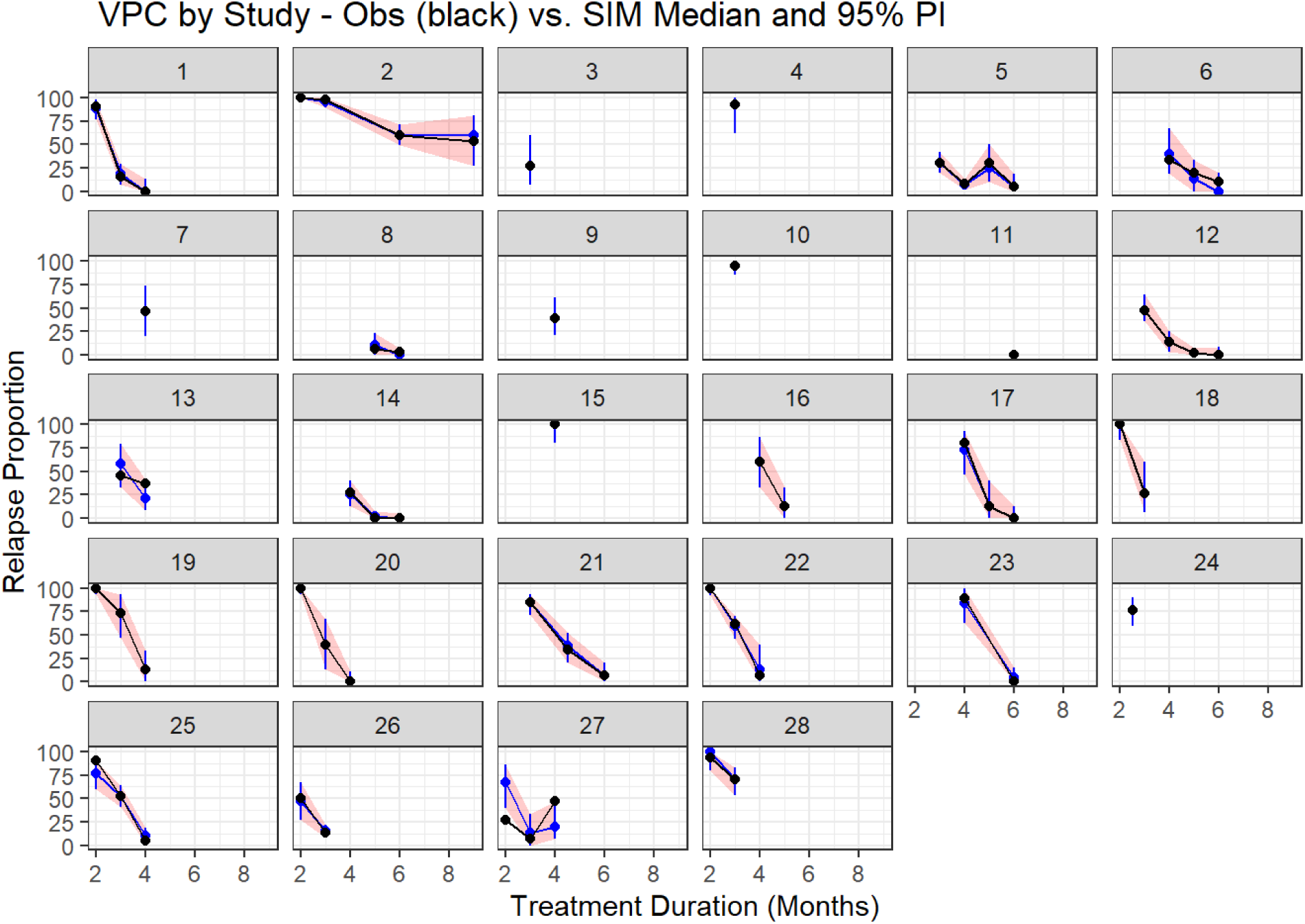
Visual predictive check (VPC) for the model used in Round 2 simulations. Red shaded regions and blue error bars represent the 90% Prediction Interval for the model. Black points and black lines represent the actual data.

**SFIG 3.**
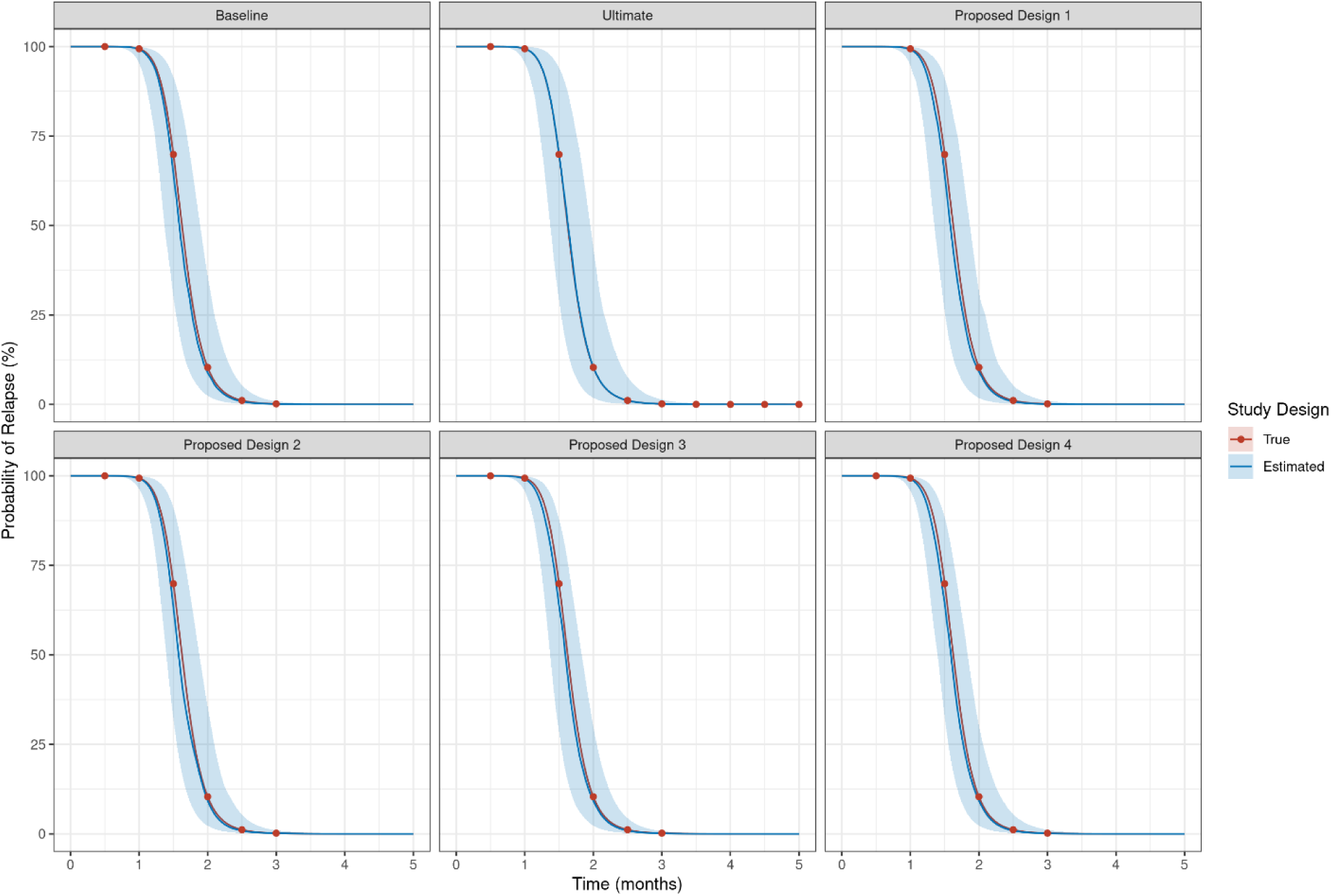
Relapse versus time profile for simulations of BPaMZ by Design for simulation round 1. Blue lines and areas represent median and 90% confidence intervals for simulations. Red lines and dots are the simulation input.

**SFIG 4.**
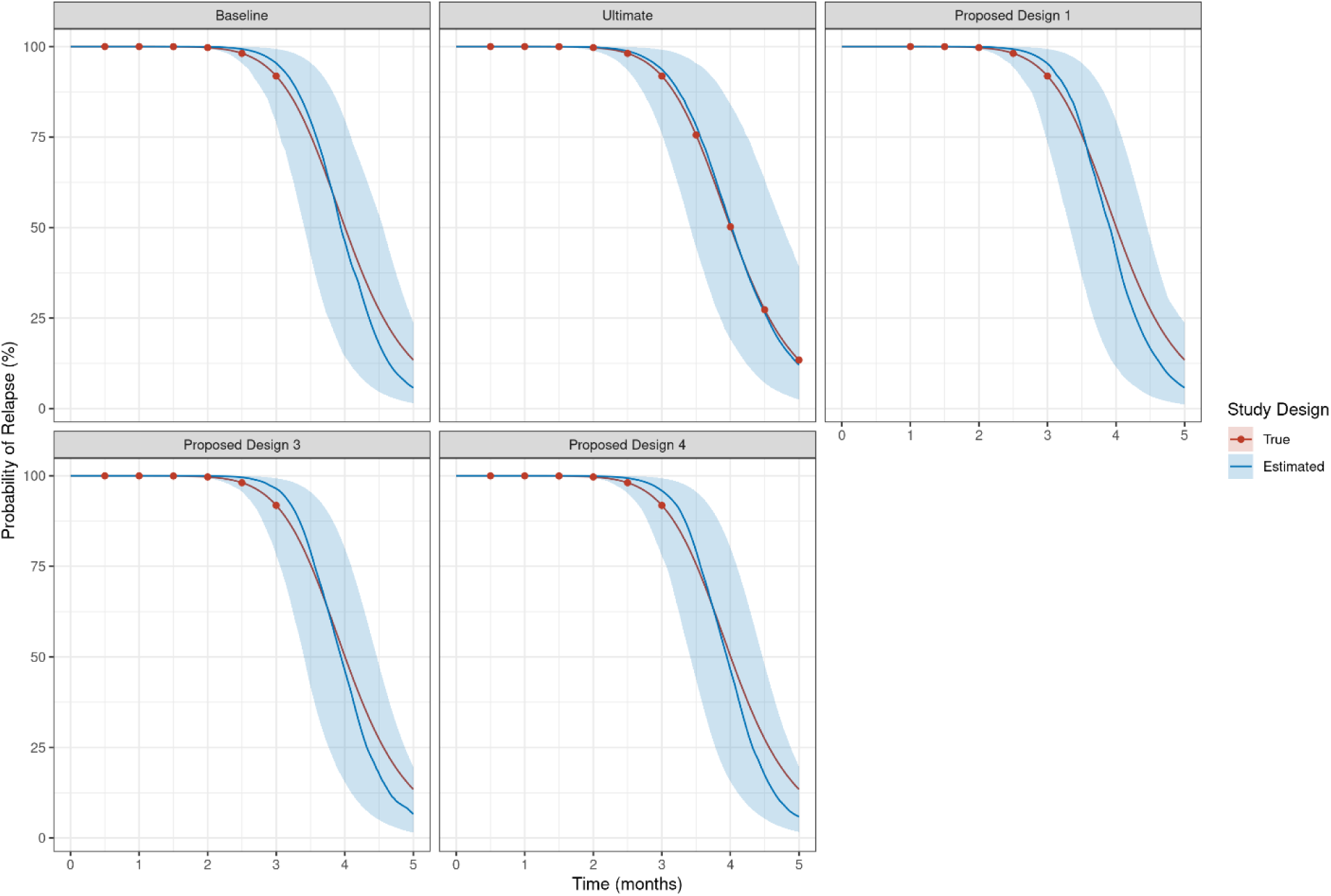
Relapse versus time profile for simulations of HRZE by Design for simulation round 1. Blue lines and areas represent median and 90% confidence intervals for simulations. Red lines and dots are the simulation input.

**SFIG 5.**
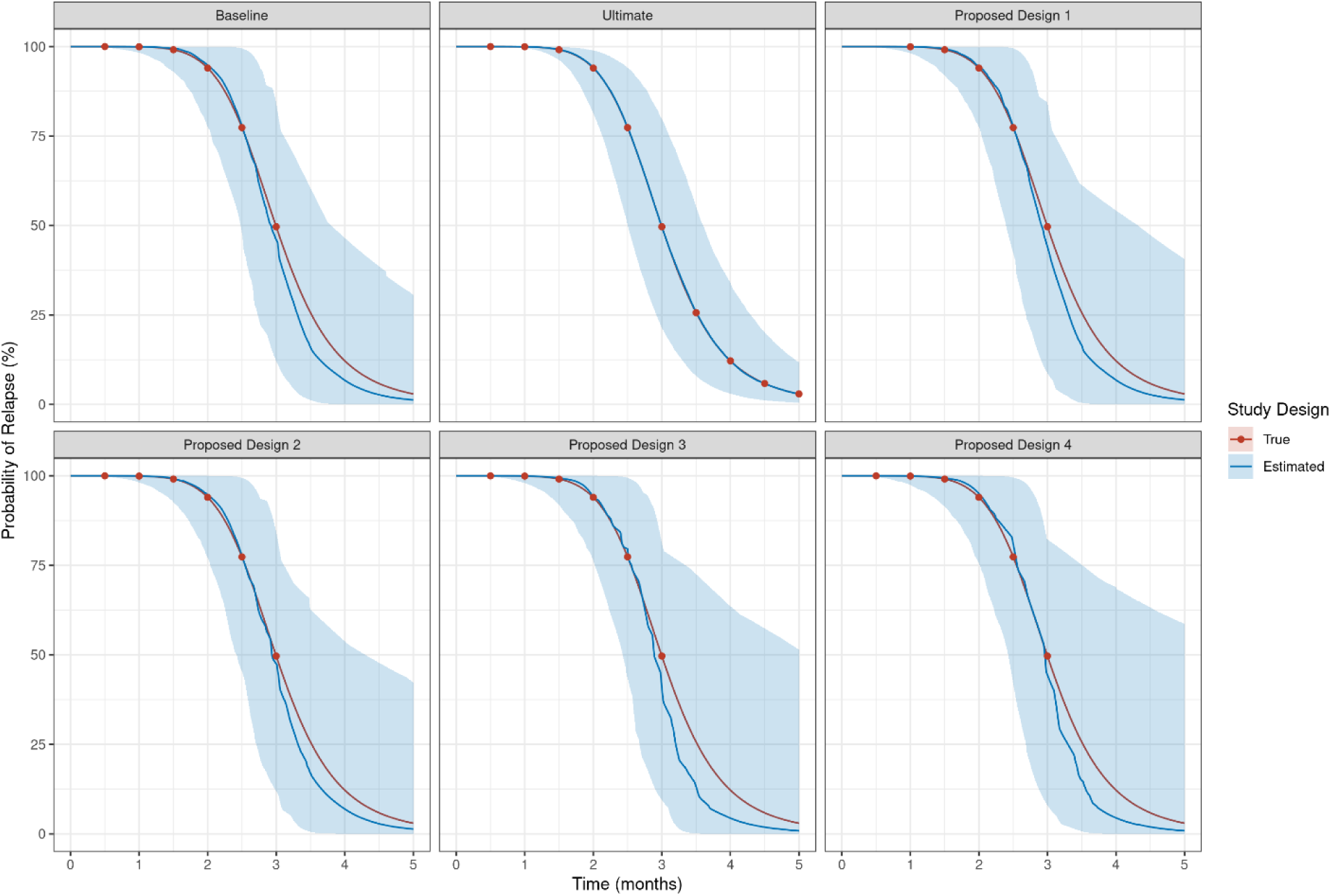
Relapse versus time profile for simulations of Regimen 1 by Design for simulation round 1. Blue lines and areas represent median and 90% confidence intervals for simulations. Red lines and dots are the simulation input.

**SFIG 6.**
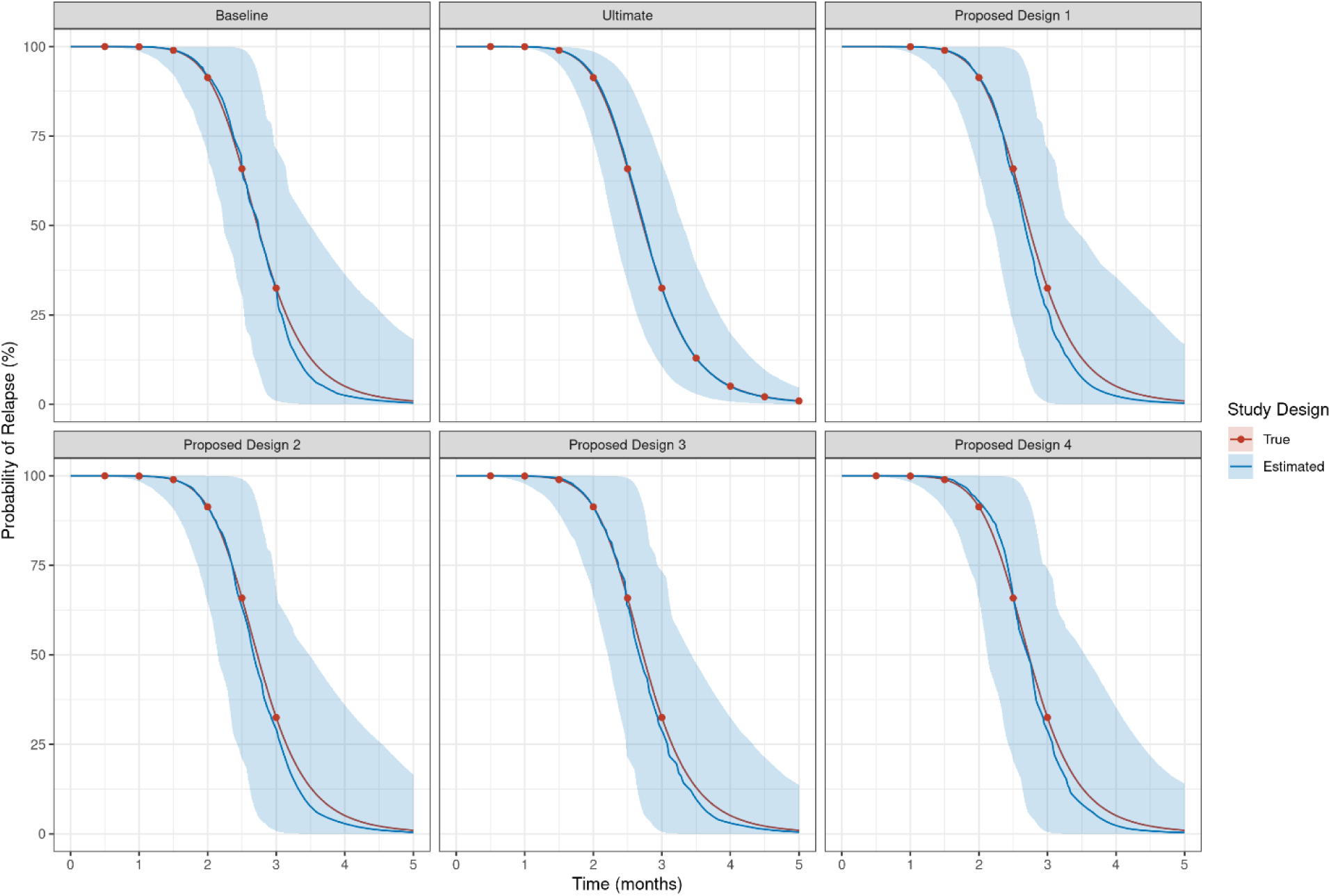
Relapse versus time profile for simulations of Regimen 2 by Design for simulation round 1. Blue lines and areas represent median and 90% confidence intervals for simulations. Red lines and dots are the simulation input.

**SFIG 7.**
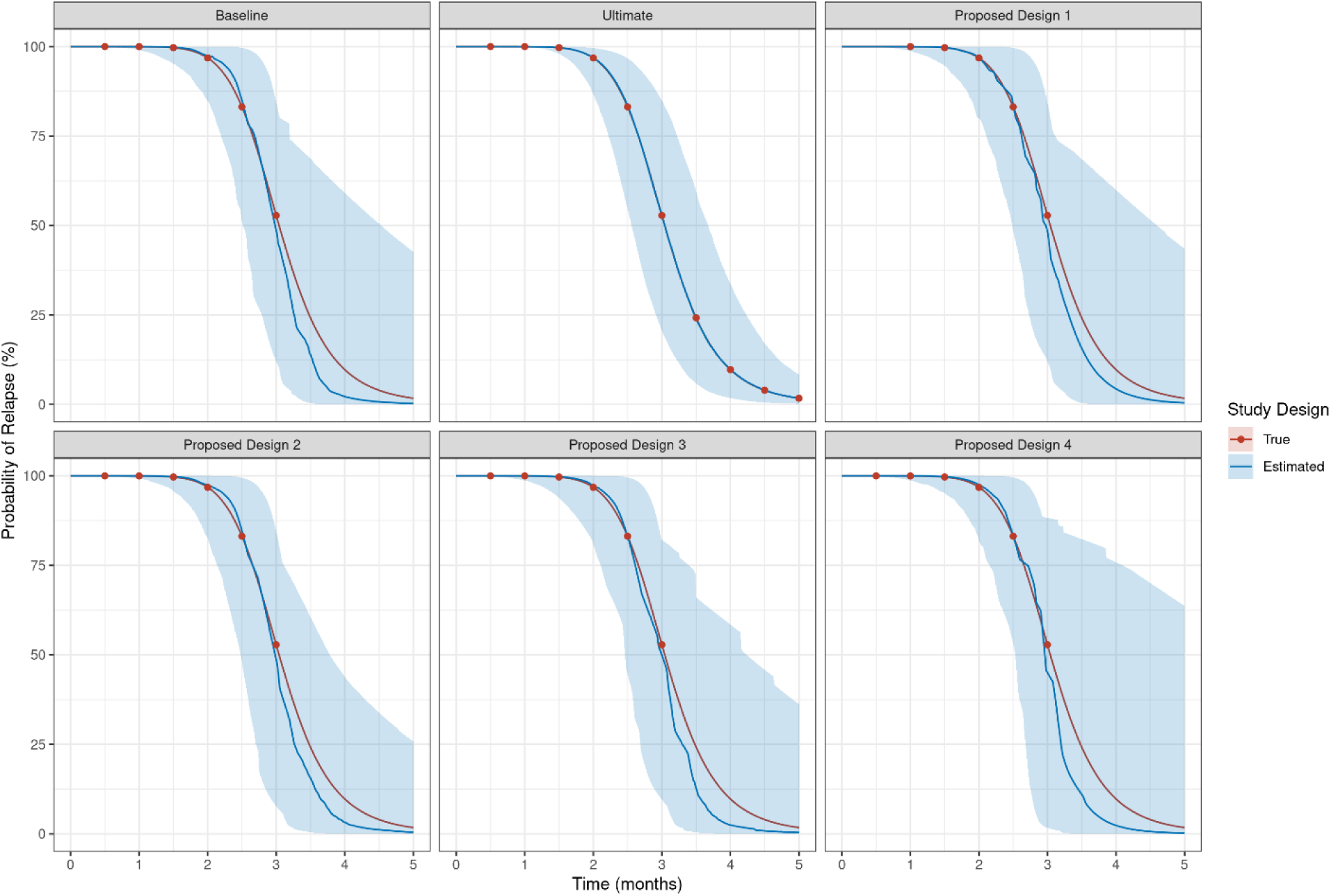
Relapse versus time profile for simulations of Regimen 3 by Design for simulation round 1. Blue lines and areas represent median and 90% confidence intervals for simulations. Red lines and dots are the simulation input.

**SFIG 8.**
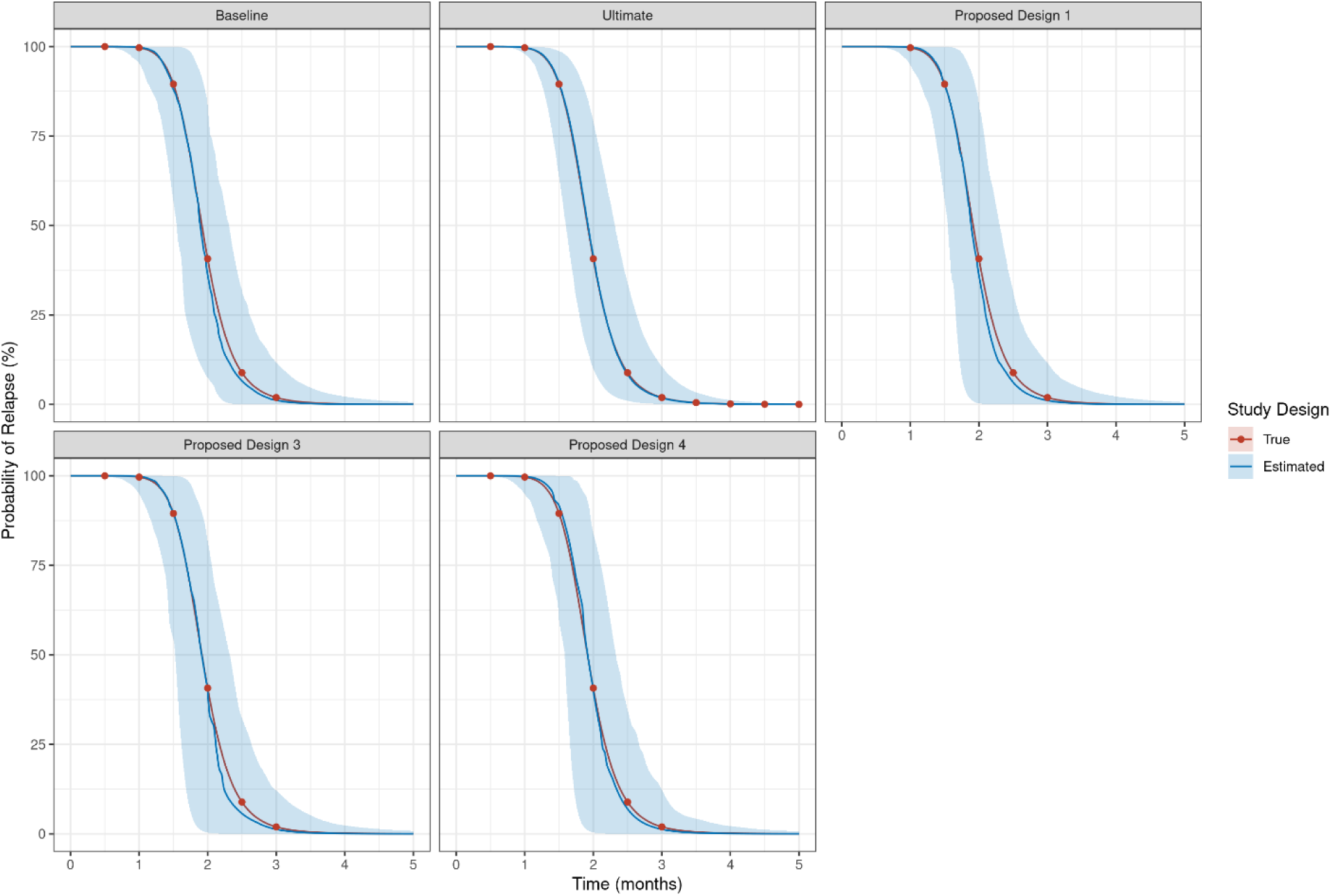
Relapse versus time profile for simulations of Regimen 4 by Design for simulation round 1. Blue lines and areas represent median and 90% confidence intervals for simulations. Red lines and dots are the simulation input.

**SFIG 9.**
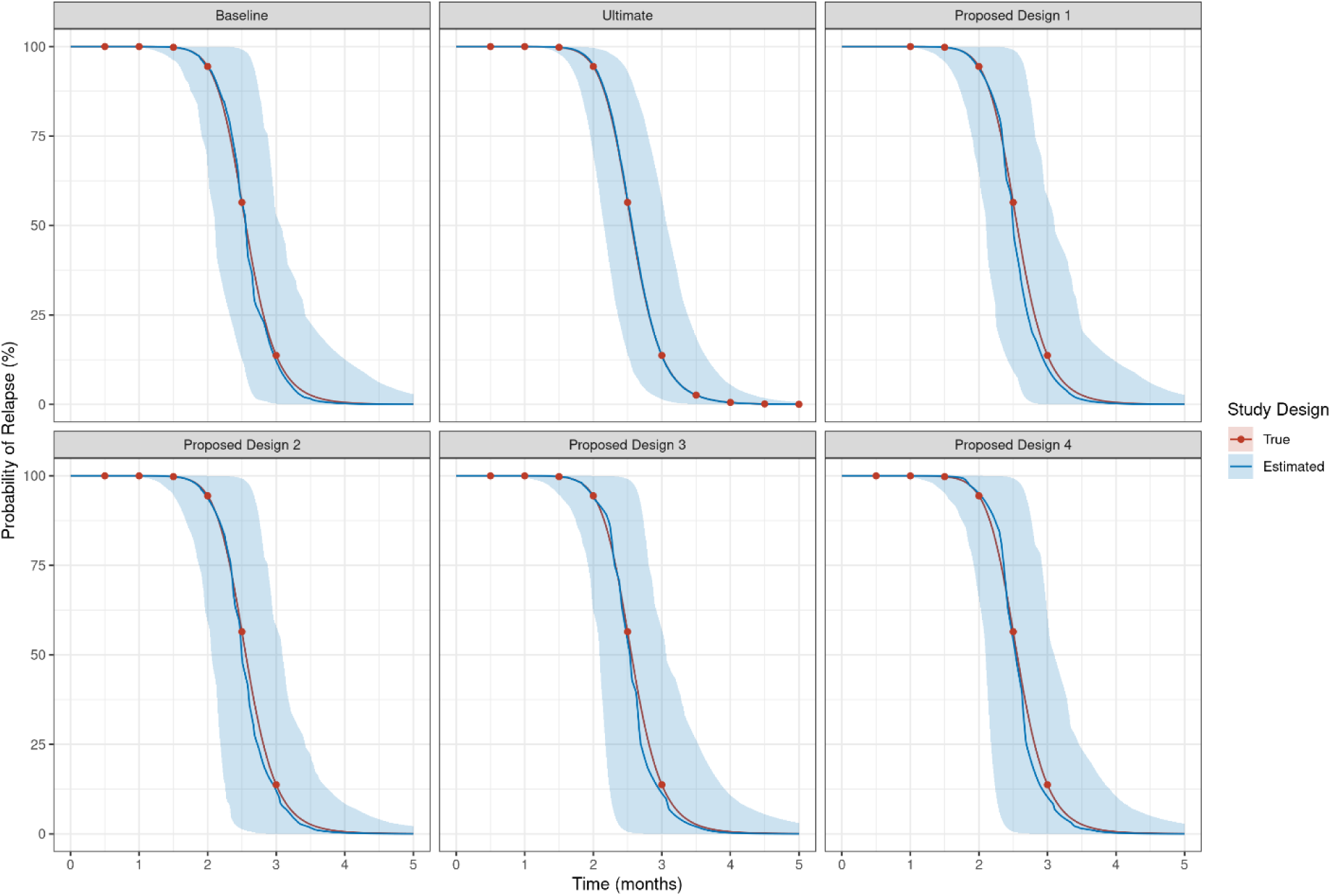
Relapse versus time profile for simulations of Regimen 5 by Design for simulation round 1. Blue lines and areas represent median and 90% confidence intervals for simulations. Red lines and dots are the simulation input.

**SFIG 10.**
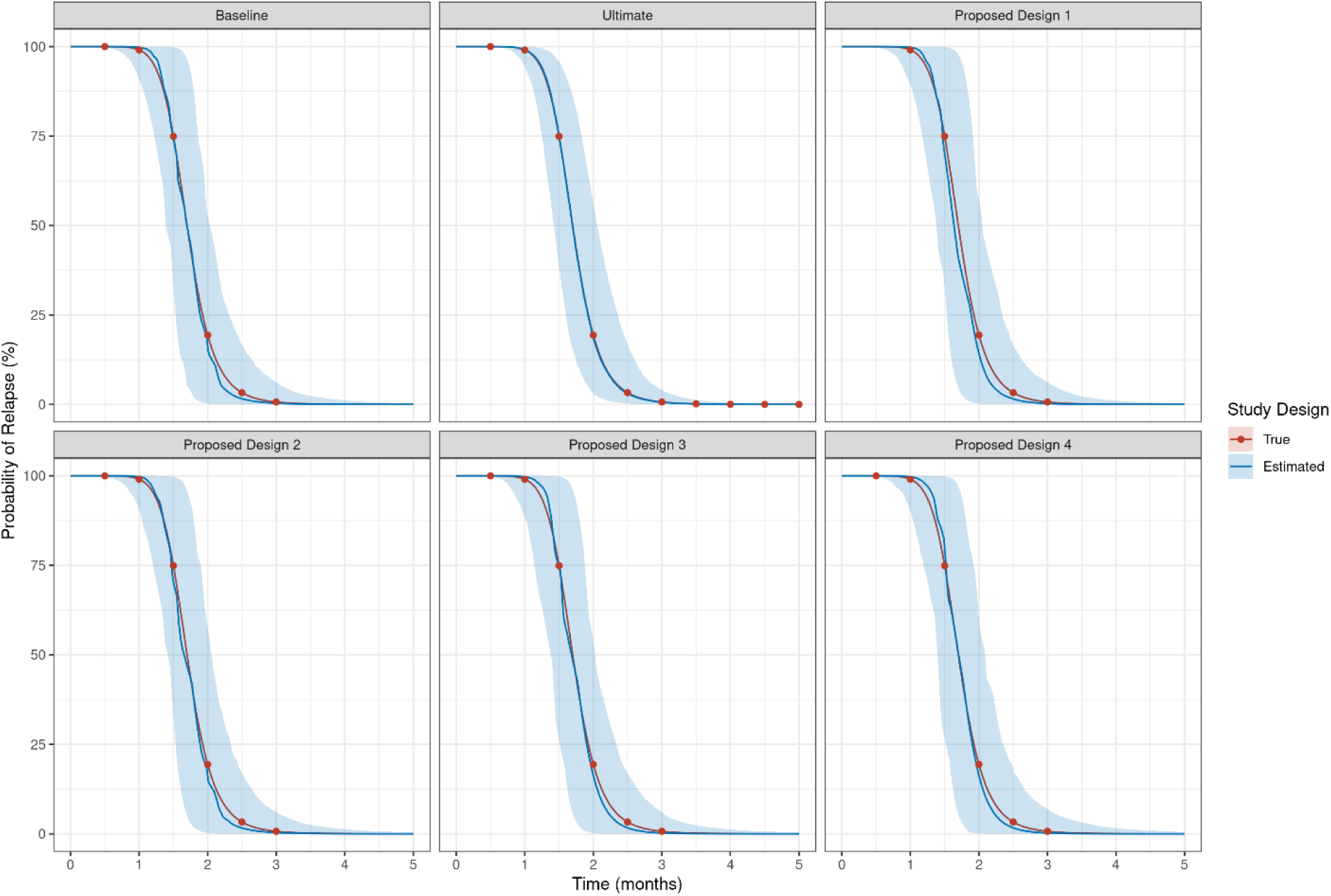
Relapse versus time profile for simulations of Regimen 6 by Design for simulation round 1. Blue lines and areas represent median and 90% confidence intervals for simulations. Red lines and dots are the simulation input.

**SFIG 11.**
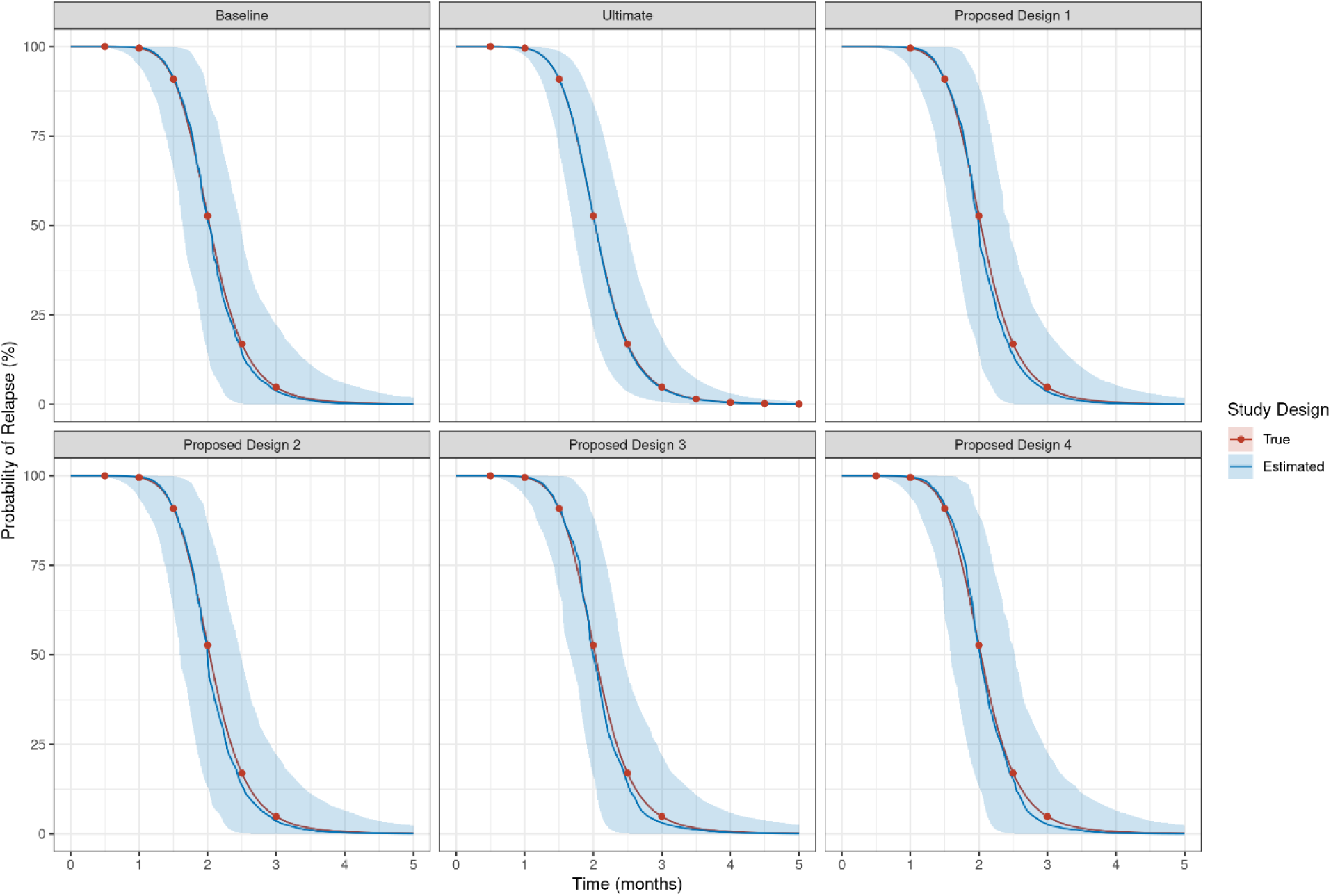
Relapse versus time profile for simulations of Regimen 7 by Design for simulation round 1. Blue lines and areas represent median and 90% confidence intervals for simulations. Red lines and dots are the simulation input.

**SFIG 12.**
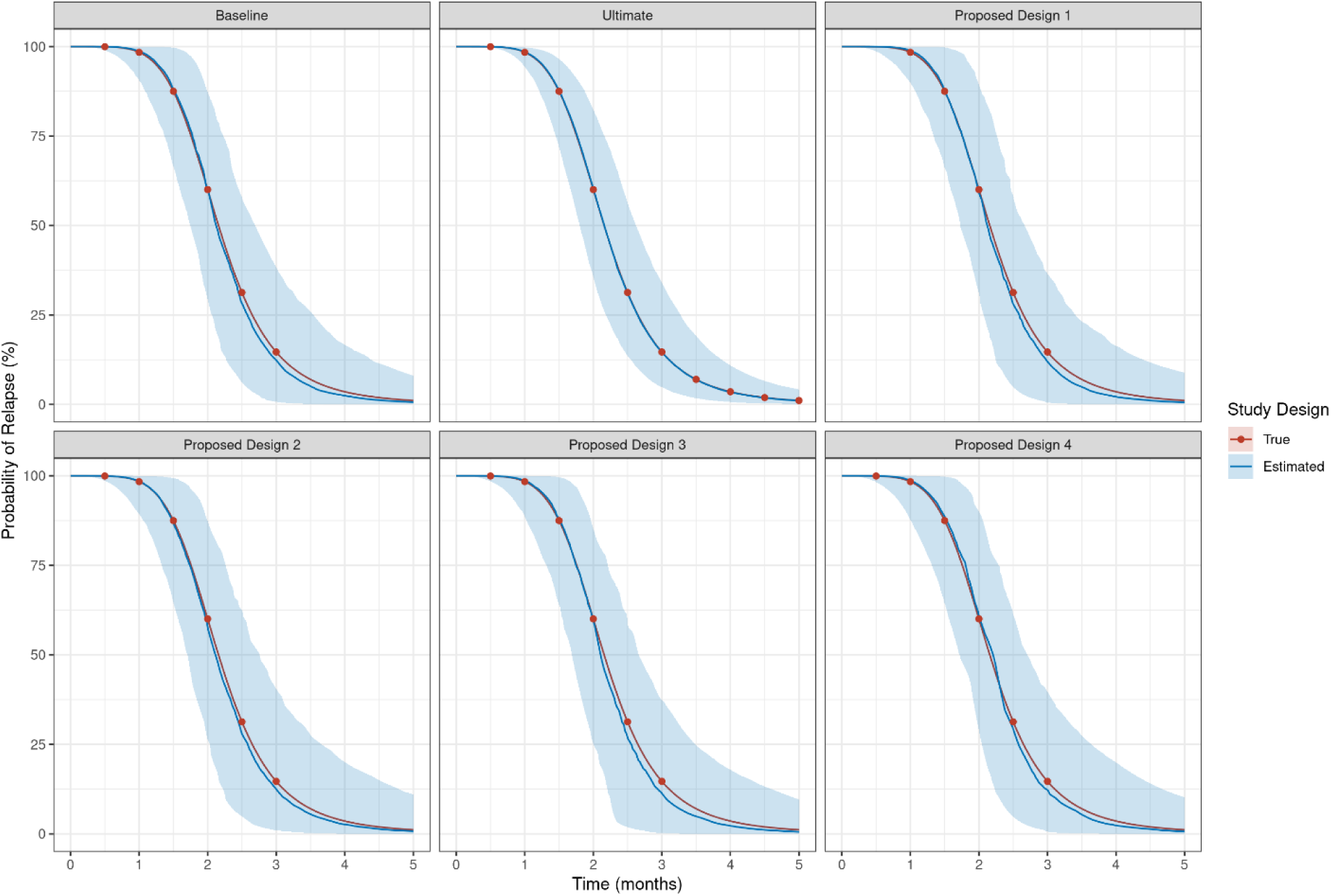
Relapse versus time profile for simulations of Regimen 8 by Design for simulation round 1. Blue lines and areas represent median and 90% confidence intervals for simulations. Red lines and dots are the simulation input.

**SFIG 13.**
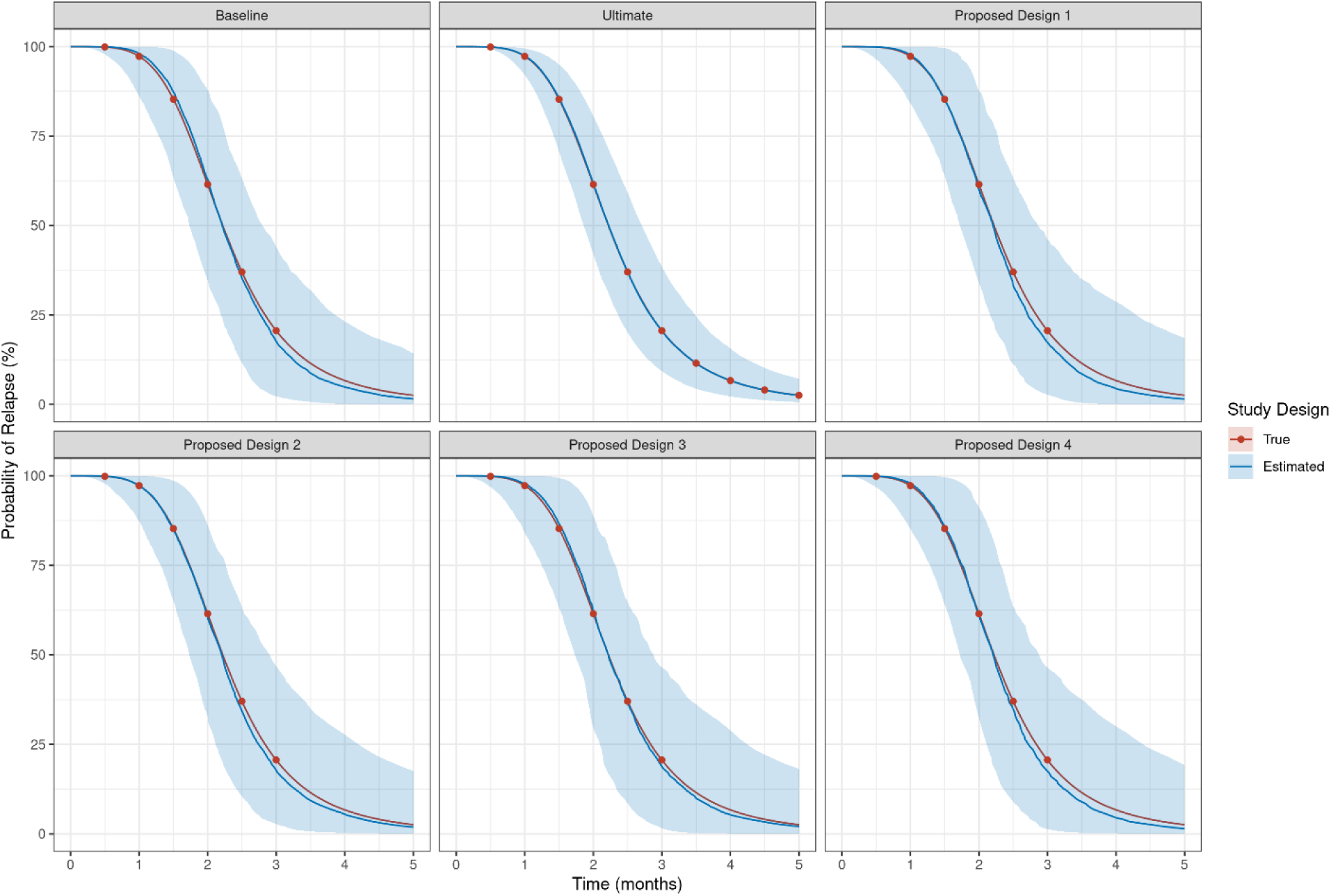
Relapse versus time profile for simulations of Regimen 9 by Design for simulation round 1. Blue lines and areas represent median and 90% confidence intervals for simulations. Red lines and dots are the simulation input.

**SFIG 14.**
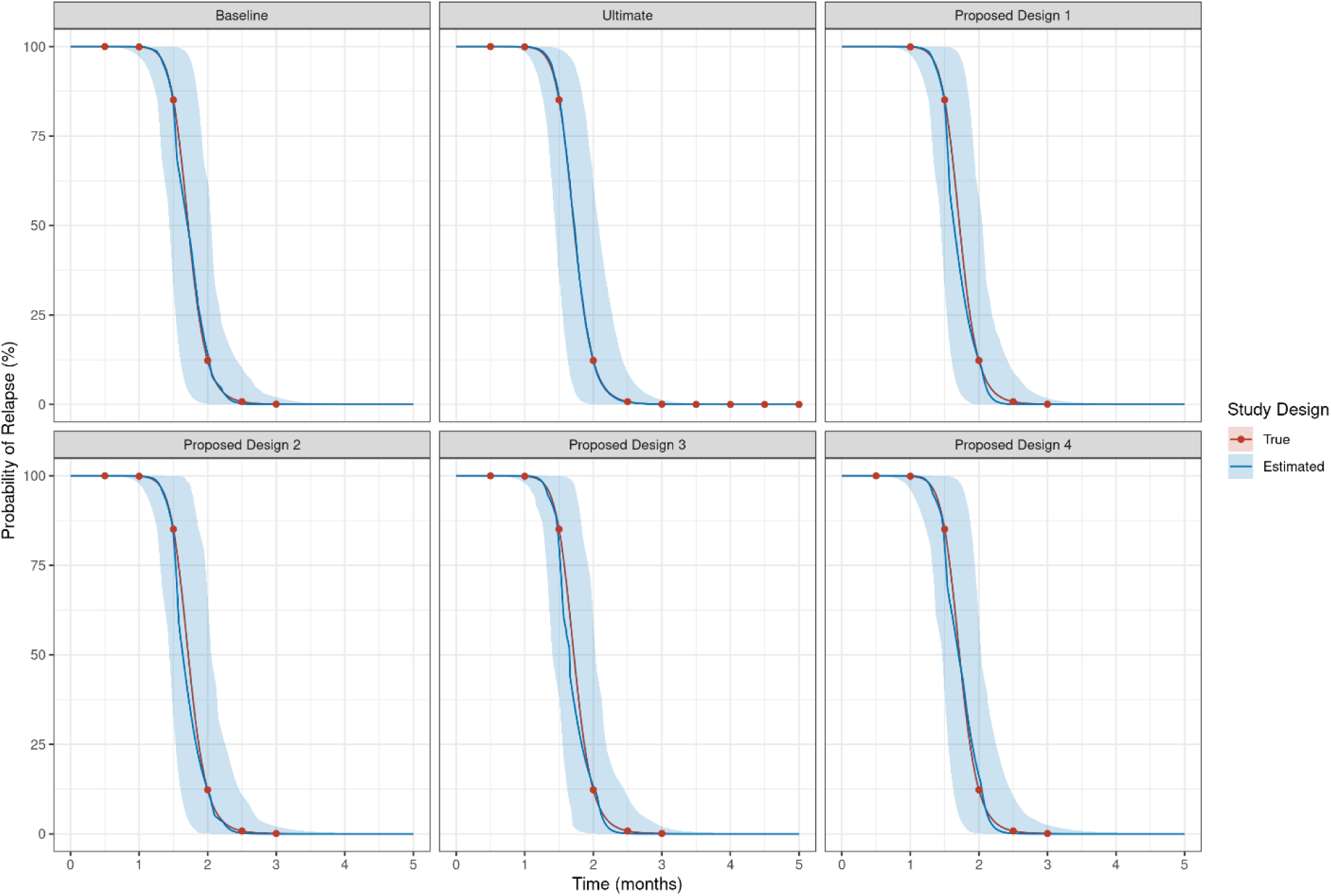
Relapse versus time profile for simulations of Regimen 10 by Design for simulation round 1. Blue lines and areas represent median and 90% confidence intervals for simulations. Red lines and dots are the simulation input.

**SFIG 15.**
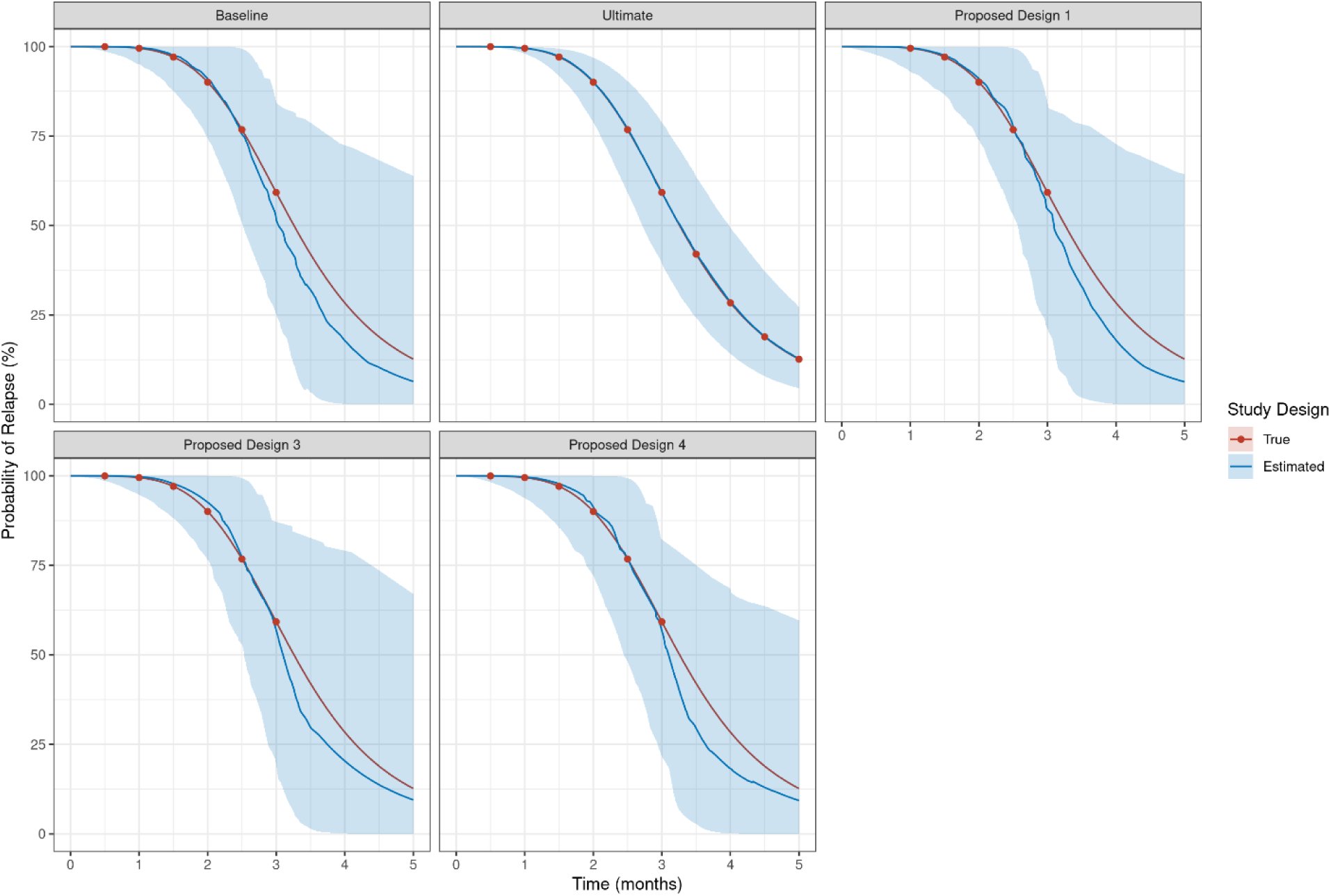
Relapse versus time profile for simulations of Regimen 11 by Design for simulation round 1. Blue lines and areas represent median and 90% confidence intervals for simulations. Red lines and dots are the simulation input.

**SFIG 16.**
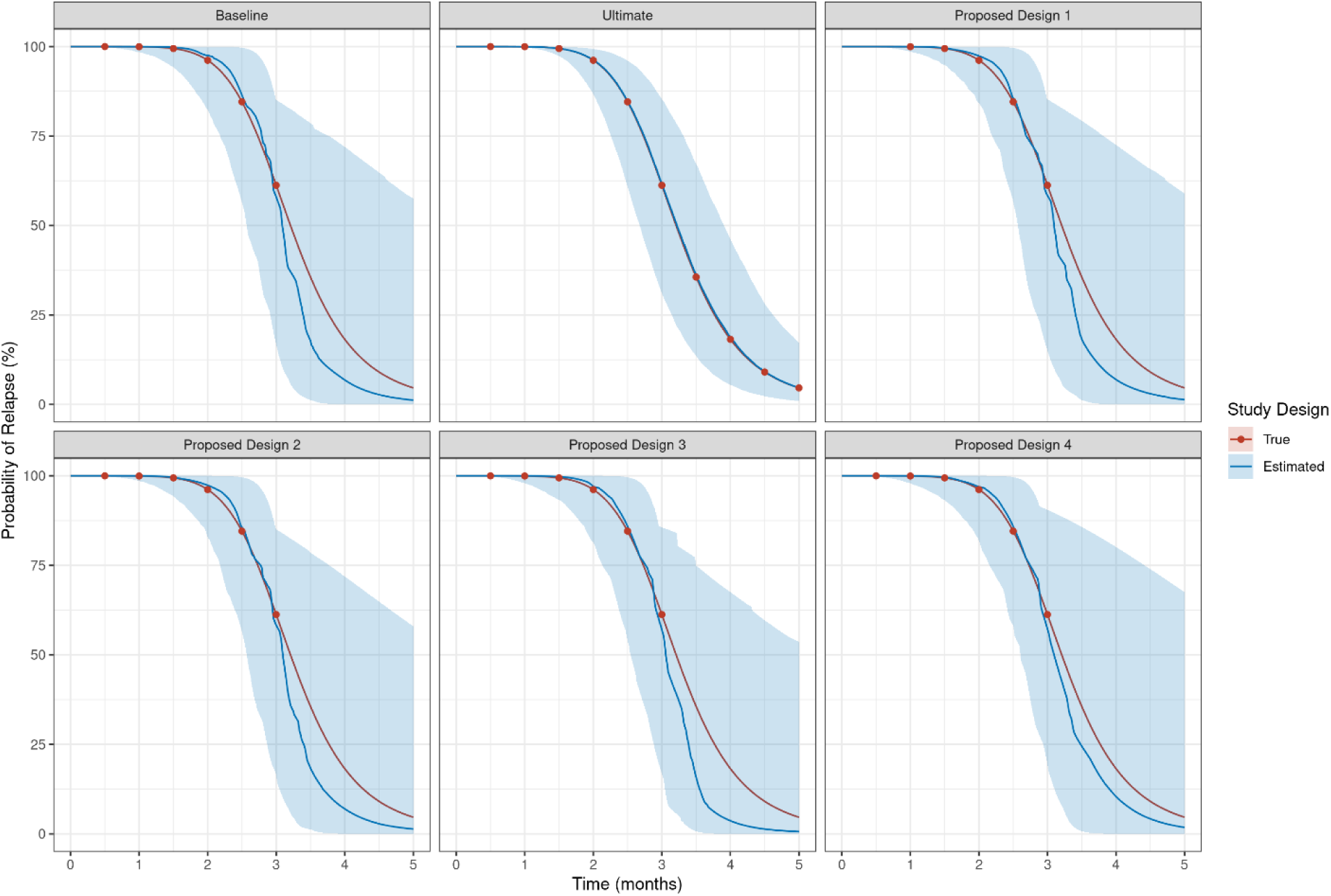
Relapse versus time profile for simulations of Regimen 12 by Design for simulation round 1. Blue lines and areas represent median and 90% confidence intervals for simulations. Red lines and dots are the simulation input.

## REFERENCES

1. World Health Organization. Global tuberculosis report 2024. 2024.

2. Carr W, Kurbatova E, Starks A, et al. Interim Guidance: 4-Month Rifapentine-Moxifloxacin Regimen for the Treatment of Drug-Susceptible Pulmonary Tuberculosis – United States, 2022. MMWR Morb Mortal Wkly Rep. 2022 Feb 25;71(8):285–289.

3. World Health Organization. Target regimen profiles for tuberculosis treatment 2023 update. 2023.

4. Franzblau SG, DeGroote MA, Cho SH, et al. Comprehensive analysis of methods used for the evaluation of compounds against Mycobacterium tuberculosis. Tuberculosis (Edinb). 2012 Nov;92(6):453–88.

5. Berg A, Clary J, Hanna D, et al. Model-Based Meta-Analysis of Relapsing Mouse Model Studies from the Critical Path to Tuberculosis Drug Regimens Initiative Database. Antimicrob Agents Chemother. 2022 Mar 15;66(3):e0179321.

6. Lenaerts AJ, Chapman PL, Orme IM. Statistical limitations to the Cornell model of latent tuberculosis infection for the study of relapse rates. Tuberculosis (Edinb). 2004;84(6):361–4.

7. Tasneen R, Betoudji F, Tyagi S, et al. Contribution of Oxazolidinones to the Efficacy of Novel Regimens Containing Bedaquiline and Pretomanid in a Mouse Model of Tuberculosis. Antimicrob Agents Chemother. 2016 Jan;60(1):270–7.

8. Sordello S BL, Tagliavini A, Federico D, Boulenc X, Pergher M, Huc CE, Metcalf D, Walter ND, Robertson GT, Clary J, Berg A, Mdluli K, Hermann D, Hanna D, Upton A. A modeling-based framework to evaluate forgiveness of TB drug combinations in a BALB/c relapsing mouse model. To Be Submitted to AAC.

9. Russell WMS, Burch RL, Hume CW. The principles of humane experimental technique. Vol. 238. Methuen London; 1959.

10. In: Weichbrod RH, Thompson GA, Norton JN, editors. Management of Animal Care and Use Programs in Research, Education, and Testing. Boca Raton (FL): CRC Press/Taylor & Francis

11. TB-Platform for the Aggregation of Preclinical Experiments Data (TB-APEX) [Internet]. 2024. Available from: https://c-path.org/tools-platforms/tb-apex/.

12. RStudio Team. RStudio: Integrated Development for R. RStudio, PBC, Boston, MA 2020.

13. Venables WN RB. Modern Applied Statistics with S. 4th ed: Springer; 2002.

14. Stan Development Team. RStan: the R interface to Stan. R package version 2.32.3. 2023.

15. Wickham H. ggplot2: Elegant Graphics for Data Analysis. Springer-Verlag New York; 2016.

